# Molecular and neuropathological determinants of neuronal dysfunction in Alzheimer’s disease

**DOI:** 10.1101/2025.10.08.681211

**Authors:** Lazaro M. Sanchez-Rodriguez, Joon Hwan Hong, Veronika Pak, Shinya Tasaki, Bernard Ng, Konstantinos Arfanakis, Pedro Rosa-Neto, David A. Bennett, Yasser Iturria-Medina

## Abstract

The biological basis of neuronal excitatory/inhibitory (E/I) imbalance in Alzheimer’s disease (AD) remains unclear. Using a comprehensive cohort with ante-mortem functional neuroimaging and post-mortem molecular data from the same participants, we mapped individual, whole-brain E/I imbalances through biophysical modeling. E/I ratios in regions supporting higher-order cognitive functions were significantly associated with cognitive performance and decline, with mediation by global neuropathological burden. We also observed a significant inverted U-shaped relationship between E/I ratios and neurofibrillary tangle severity, peaking at the limbic stage (Braak III-IV) in 14 brain areas, including the bilateral hippocampus and superior frontal gyrus. In addition, we identified 89 genes and 101 proteins that predict regional E/I ratios, with pathways related to synaptic signaling and immune response overrepresented. The generalizability of these molecular predictors was confirmed in two independent cohorts, achieving good classification performance for neuropathology severity and AD dementia. Lastly, the estimated E/I imbalances in AD aligned with whole-brain distributions of microglia and oligodendrocyte precursor cells, suggesting that spatial cellular organization contributes to vulnerability to neuronal dysfunction. Overall, this study provides critical insights into the cellular, molecular, and neuropathological signatures of circuit-level dysfunction in AD.

## Introduction

The balance between neuronal excitatory (E) and inhibitory (I) signaling is critical for stable information processing in the human cortex^1^. It has been proposed that disruption of this balance parallels progression of Alzheimer’s disease (AD)^2^. Early evidence from animal models^3–5^, post-mortem surrogate cell analyses^6,7^, and indirect electrophysiological assessments^8,9^ suggests a pro-excitatory shift in brain activity in AD, linked to the aggregation of amyloid-beta (Aβ) and tau proteins^4,10–13^. These neuropathological changes contribute to synaptic dysfunction, impaired network activity, and ultimately to cognitive decline^2,14^. Dysregulation of the E/I ratio has thus become a fundamental aspect bridging AD’s neuropathological features to their clinical manifestations, positioning it as a potential phenotypic readout for AD therapies^2^. However, as in-vivo assessments of E/I ratios are challenging, the molecular factors driving E/I imbalances in human AD remain poorly understood ^7–9^.

Numerous computational approaches have been devised to improve our understanding of the relation of E/I ratios to human behavior. Calculating the aperiodic component, or "1/f slope", of the electrophysiological (EEG/MEG) power spectrum has become the gold-standard proxy for measuring E/I balance^15,16^. This metric is based on computational simulations of a single excitatory and a single inhibitory neuronal population interacting under specific conditions^16–18^. In prior work, we used^19,20^ resting-state functional MRI (rs-fMRI)^21,22^, computational models^23–25^, and whole-brain gene expression data from neurotypical brains^26,27^ to investigate neuronal activity in AD. We observed increased excitatory neuronal activity, estimated from rs-fMRI, associated with Aβ positivity and severe tau deposition in limbic and cortical brain regions^20^. These functional metrics predicted changes in brain structure, cognitive decline, and brain-wide involvement of cellular signaling, architectural and inflammatory molecular pathways^19,20^. However, these findings were limited by the use of gene expression data from a neurotypical brain template, rather than directly from AD-affected tissue of matched individuals. Other studies have identified individual multilevel molecular markers of AD progression^28–30^, but they failed to incorporate measures of neuronal dynamics.

To obtain a more precise molecular characterization of the E/I imbalance in AD, we leverage a unique dataset from the Religious Order Study and the Rush Memory and Aging Project (ROSMAP), which includes ante-mortem resting-state fMRI and cognitive data, along with post-mortem neuropathological assessments, in addition to gene expression and protein abundance from AD-relevant brain regions^31,32^. The objectives of this study are three-fold. First, we generated individual E/I ratio maps and characterized the role of neuronal E/I imbalance in AD clinical and pathological progression. Second, we identified transcriptomic and proteomic signatures that explain E/I imbalances in the AD brain, utilizing post-mortem molecular profiles from the same participants. We evaluated the generalizability of the identified molecular markers for predicting neuropathological severity and AD dementia in two independent cohorts. Thirdly, we tested whether the spatial distributions of six major cell types in neurotypical brains are associated with regional vulnerability to AD-related E/I imbalances.

## Results

### Personalized biophysical analysis to map E/I imbalance in Alzheimer’s disease

We developed personalized neural mass models to estimate key biophysical parameters, including local neuronal excitability and inter-regional coupling strength, from ante-mortem resting-state fMRI (rs-fMRI) data of 156 participants from the Religious Orders Study and Rush Memory and Aging Project (ROSMAP, see Supplementary Table 1)^31–33^. Participants were categorized as no cognitive impairment (NCI), mild cognitive impairment (MCI), or AD dementia, based on their most likely clinical diagnosis at the time of death^34–36^. Neuropsychological evaluations included a global measure of cognition and summary measures of five domains of cognitive function (episodic memory, visuospatial ability, perceptual speed, semantic memory, and working memory) and their rates of changes over time^37,38^. A global AD pathology burden was obtained from standardized counts of neuritic plaques (NPs), diffuse plaques, and neurofibrillary tangles (NFTs) across five regions: midfrontal cortex, midtemporal cortex, inferior parietal cortex, entorhinal cortex, and hippocampus (CA1)^39,40^. NPs and NFTs were also summarized as semiquantitative estimates using the Consortium to Establish a Registry for Alzheimer’s Disease (CERAD) criteria and Braak staging^32,41,42^. Other neuropathologic assessments included Lewy body disease, TDP-43 pathology, hippocampal sclerosis, arteriolosclerosis, cerebral atherosclerosis, presence of chronic macroinfarctions and microinfarctions^43–47^. Brain-derived omics data included bulk RNA-seq (12,905 genes) and protein abundance (7,788 proteins) from the superior frontal gyrus (SFG) and inferior temporal gyrus (ITG) for a subset of 87 matched participants (Supplementary Table 2).

The biophysical parameters of interest were inferred by maximizing the similarity between the model-generated and real rs-fMRI signals, specifically focusing on the regional amplitude of low-frequency fluctuations (ALFF, 0.01–0.08 Hz). ALFF is a well-established marker of spontaneous neuronal activity sensitive to AD progression^48–50^. For each participant, ALFF is computed at the voxel-level as the average power within the frequency band of interest and subsequently averaged at the region-level^51^. We used a fast machine learning algorithm coupled with surrogate optimization techniques to estimate parameters (Supplementary Figure 1; *Methods, Estimating neuronal activity alterations*). Through the individual biophysical parameters, we reconstructed underlying excitatory and inhibitory signals and estimated regional E/I ratios for each subject across the brain^20,52–55^, thereby capturing emergent functional properties of the brain (Figure 1).

**Figure 1.**
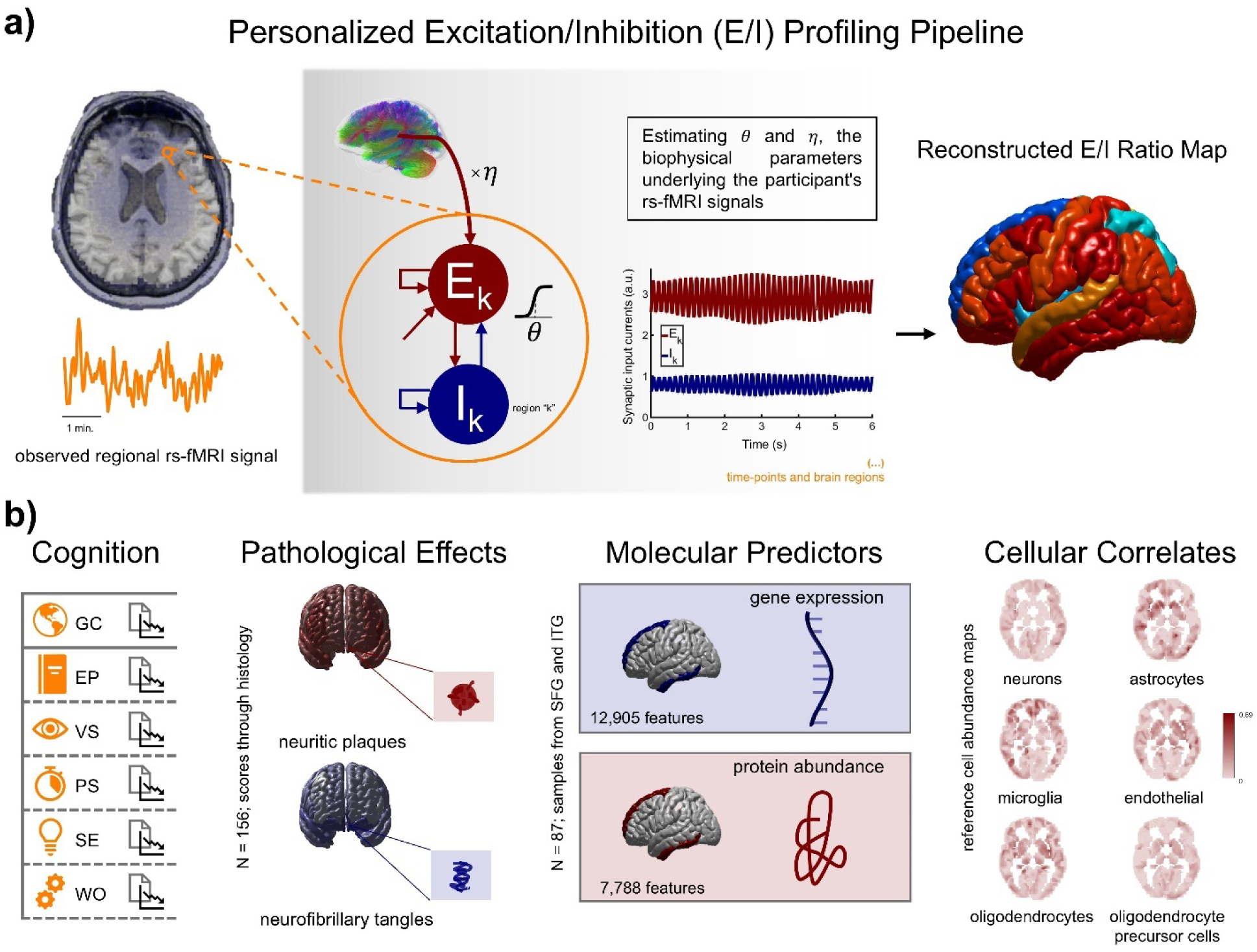
Overview of the computational pipeline for multi-scale integration. **a**) A neural mass model with excitatory (E) and inhibitory (I) interactions is used to estimate biophysical parameters from the participant’s rs-fMRI signals. Key parameters—firing threshold (θ) and coupling strength (η)—regulate local neuronal excitability and inter-regional communication. An optimization algorithm identifies the θ and η values that best match the participant’s rs-fMRI markers (regional amplitudes of low-frequency fluctuations, ALFF). Whole-brain E/I ratio maps are generated from the optimal θ and η values. **b**) Relationships between the E/I ratio maps and disease-relevant data were investigated, including cognitive performance and its temporal decline, neuropathological evaluations (e.g., measures of neuritic plaques and neurofibrillary tangles), post-mortem bulk gene expression and protein abundance data from the superior frontal gyrus (SFG) and inferior temporal gyrus (ITG), and template cell abundance maps for six major brain cell types.

Subsequently, we examined the relationships between the obtained subject-specific E/I ratio maps and a range of cognitive, neuropathological, molecular and cellular measures. All analyses were statistically adjusted for age, sex, education years and MRI acquisition protocol. In addition, for the participants with available post-mortem gene expression and protein abundance data, we applied multivariate predictive modeling techniques to identify brain transcriptomic and proteomic markers predicting the E/I ratios in these regions. We also assessed whether this set of molecular predictors generalized to independent cohorts comprising different participants, brain regions, and experimental conditions.

### Regional E/I ratios predict cognitive performance

We proceeded to test whether the participants’ estimated regional E/I ratios (across the 104 brain regions) explain individual multi-domain cognitive performance, as well as their temporal rates of decline. For each cognitive variable, we used elastic net regression (10-fold cross-validation), including sex, age, education, and MRI protocol type as covariates. The significance of regional coefficients was assessed through 1,000 permutations per cognitive variable, followed by max-T correction for family-wise error rate (FWER, p < 0.05). Elastic net regression removes redundant predictors that do not contribute significantly to explaining the response variables, thereby mitigating multicollinearity, improving interpretability, and reducing model complexity^56^.

Figure 2 summarizes the regional coefficients for each cognitive score, highlighting predictors that passed FWER correction. These results reveal distinct spatial patterns across cognitive domains and their decline. Notably, the E/I ratios at the right lingual gyrus and right insula emerged as consistent predictors of both global cognition and episodic memory, further highlighting the role of these regions in high domain cognitive functions^57,58^. The left hippocampus, a structure with a well-established role in memory encoding and retrieval^58,59^, was also significantly associated with both scores. Processing speed was predicted by E/I at the left thalamus, while the right insula also related to semantic memory. For cognitive decline, we found associations between the lingual and transverse temporal gyri and slope of global cognition, and between the left caudate and slope of working memory.

**Figure 2.**
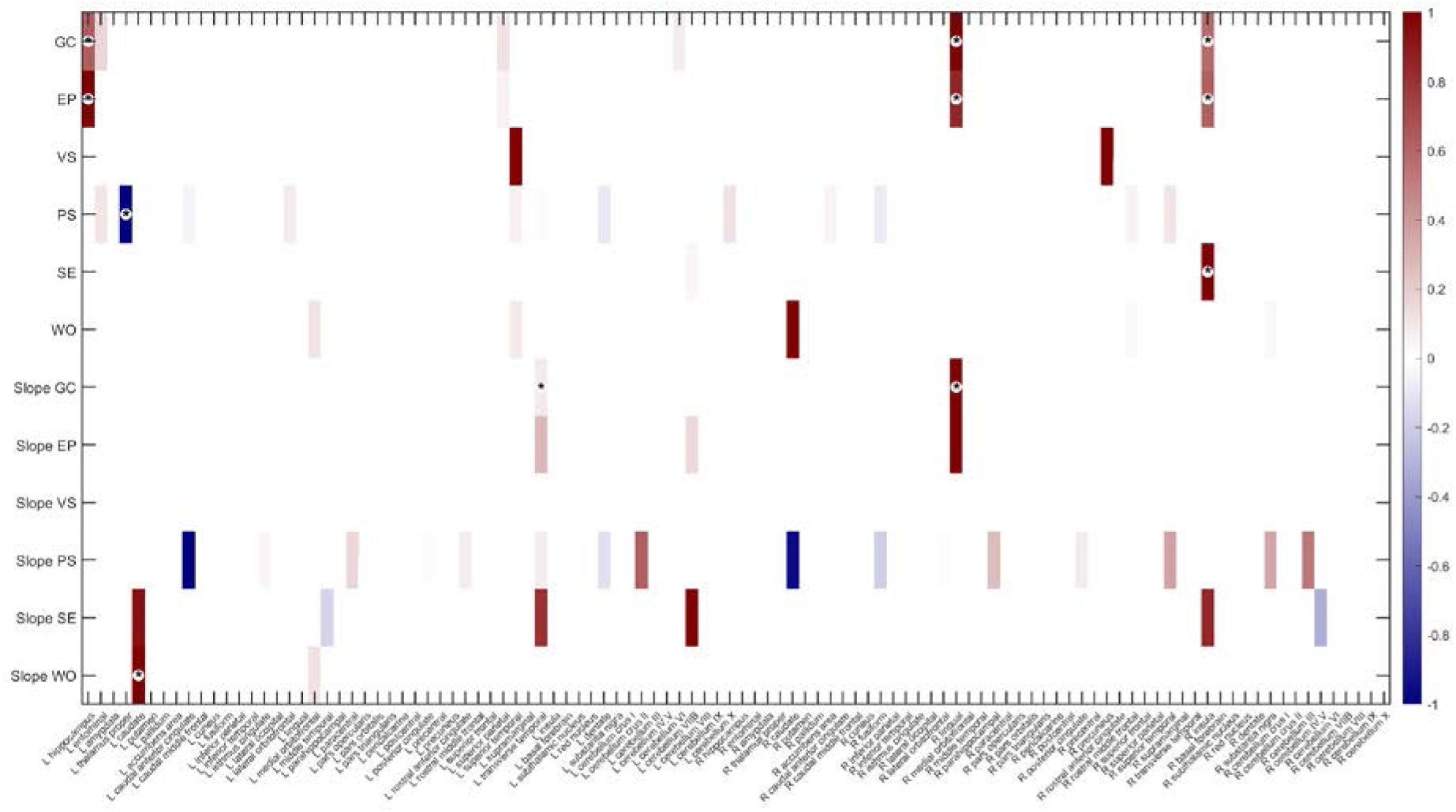
Regional E/I ratio relationships with cognitive scores across 104 ROIs. Elastic net regression coefficients are shown for predictors selected for each cognitive score, with positive (maroon) and negative (navy) relationships. ROIs marked with asterisks indicate regions surviving FWER correction at p < 0.05 based on permutation testing. Coefficients have been normalized to the [–1, 1] range for visualization purposes (see Supplementary Table 3 for full coefficient values and p-values). GC, global cognition; EP, episodic memory; VS, visuospatial ability; PS, perceptual speed; SE, semantic memory; WO, working memory. No predictors were selected for the visuospatial slope score.

While multiple regions were selected by the elastic net models for each score (see Supplementary Table 3), the FWER-corrected findings point to a consistent set of E/I ratio features—particularly in the medial temporal lobe, insula, thalamus, and striatum—as being most robustly linked to individual differences in cognition and cognitive change. Among these selected predictors, most relationships were positive, indicating that lower E/I ratios in these regions are associated with poorer cognitive performance. However, some inverse effects were also observed (e.g., in the left thalamus). Overall, these findings support an additive effect of E/I ratios across AD-related regions on multiple cognitive domains.

### Regional E/I ratios strongly associate with neurofibrillary tangle density

Next, we examined how the estimated regional E/I ratios relate to AD’s neuropathological hallmarks. Specifically, we tested the associations between E/I ratios and a global quantitative summary of AD pathology (Global AD)^39,40^, derived from counts of neuritic plaques, diffuse plaques, and NFTs in five regions of interest, as well as overall measures of Aβ and tangle burden. To this end, we used partial correlations and permutation-based tests between each pair of variables of interest, controlling for age, sex, years of education, and MRI protocol type. Notably, we observed significant negative associations between Global AD pathology and E/I ratios in the left hippocampus, left superior parietal cortex, bilateral insulae, right superior frontal cortex, right lingual gyrus, and right caudate nucleus (all p < 0.05, FWER-corrected via max-T permutation using Fisher Z-values across 1,000 iterations; Figure 3a, Supplementary Table 4). Similarly, NFT density showed significant negative associations with E/I ratios in a broader set of regions, including bilateral superior frontal gyrus, hippocampus, caudate, putamen, insula, and thalamus, as well as the left superior parietal cortex and cerebellar region 7b. These findings suggest a consistent pattern in which higher pathology is linked to lower E/I ratios across both cortical and subcortical structures.

**Figure 3.**
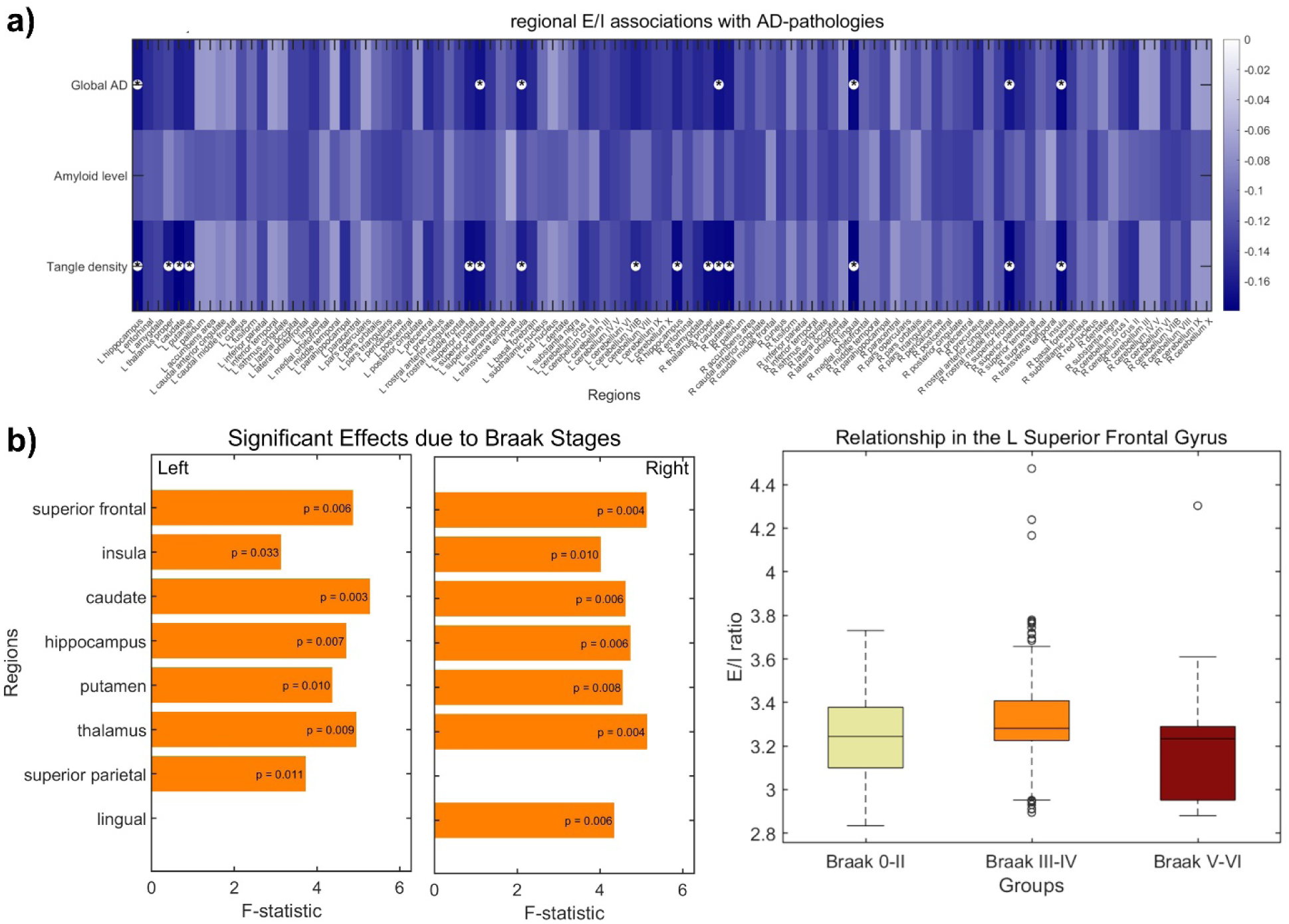
Associations between regional E/I ratios neuropathologies. **a**) Pearson correlation values for the relationships between regional E/I ratios and global AD pathology (Global AD), amyloid (Aβ), and neurofibrillary tangle (NFT) burden across 104 brain regions. Analyses were adjusted for age, sex, years of education, and MRI protocol. Asterisks indicate regions with significant associations (p < 0.05, FWER-corrected via max-T permutation). **b**) Effects of NFT severity (Braak stage) on regional E/I ratios. Left: regions showing significant effects (p < 0.05, FWER-corrected) based on multi-way ANOVA with permutation testing, with corresponding F-values shown separately for the left (L) and right (R) hemispheres. Right: box-and-whisker plot illustrating the non-linear association between Braak stage and E/I ratio in the left superior frontal gyrus. Boxes indicate interquartile range, central lines show medians, whiskers extend to the most extreme non-outlier values, and individual outliers are shown as circles.

**Figure 4.**
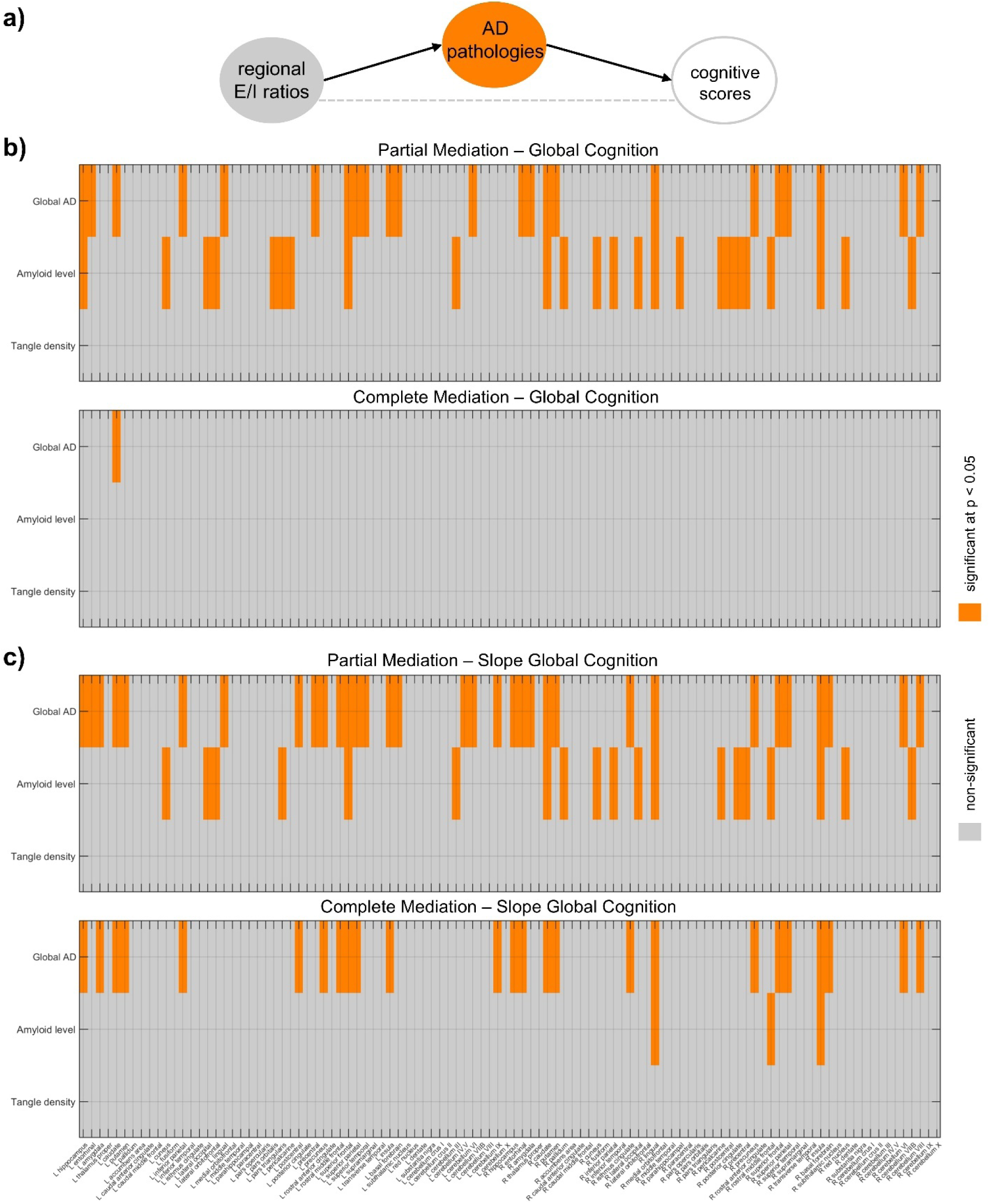
Mediation analysis of the relationship between regional E/I ratios and cognitive scores, with AD pathologies as mediators. **a**) Schematic representation of the mediation model. **b**) Associations between regional E/I ratios and baseline global cognition mediated by AD pathologies. **c**) Associations between regional E/I ratios and the slope of global cognition mediated by AD pathologies. Orange boxes indicate significant effects (p < 0.05).

To further understand these relationships and identify key disease stages, we performed a multi-way ANOVA testing whether regional E/I ratios differ significantly across levels of neuropathological burden. We included both AD-related semi-quantitative staging variables (CERAD scores and Braak stages) and non-AD-specific pathologies, including Lewy body disease, TDP-43 pathology, hippocampal sclerosis, arteriolosclerosis, cerebral amyloid angiopathy, atherosclerosis, and the presence of cortical infarcts. Demographic and scanning covariates (age, sex, years of education, MRI protocol) were also included. To control for multiple comparisons across regions and account for imbalanced group sizes, we applied a permutation-based max-F correction using 1,000 iterations. While no effects of the participants’ CERAD scores were found, a significant inverted U-shaped pattern emerged with Braak stages of NFT severity (Figure 3b): E/I ratios were low at Braak stages 0-II (absence of pathology or pathology mainly confined to the entorhinal region), increased at Braak stages III-IV (the limbic stage), and decreased again at Braak stages V-VI (the neocortical stage). For illustrative purposes, the pattern at the left superior frontal gyrus is shown in the figure, while similar trajectories in several consistently affected regions—including the bilateral superior frontal gyrus, caudate, hippocampus, putamen, thalamus, insula, the left superior parietal cortex, and the right lingual gyrus—are presented in Supplementary Figure 2. Among non-AD pathologies, hippocampal sclerosis and atherosclerosis showed significant associations with E/I ratios across distinct sets of regions (11 and 45 regions, respectively), further supporting the sensitivity of E/I balance to diverse neuropathological processes. Effects of hippocampal sclerosis on E/I ratios were primarily observed in subcortical and medial temporal areas, whereas atherosclerosis was associated with widespread alterations spanning frontal, temporal, parietal, and subcortical regions, as well as several cerebellar structures (Supplementary Table 5).

The observed non-linear trajectory of E/I balance across Braak stages highlights a possible inflection point in AD progression. Increasing evidence suggests that Braak stages III–IV represent a critical transition in terms of cognitive decline^64,65^ and neuropathological interactions^20,21,66^. Our findings suggest that this may also be a turning point in the dynamics of regional E/I balance.

### AD pathologies as mediators of the relationship between regional E/I ratios and cognitive performance

We conducted a mediation analysis^67–69^ to examine the potential mediating role of AD pathology in the association between regional E/I ratios and cognitive performance, as well as its temporal decline. This was followed by a reversed analysis, considering E/I ratios as mediators in the association between AD pathology and cognitive performance and decline. Specifically, we considered Global AD, average Aβ, and NFTs as pathological variables of interest, and global cognition and its annual slope as cognitive outcomes. For each case (i.e., with E/I as the main independent variable and pathology as the mediator, or alternatively, with E/I as the mediator of pathology), we tested two models: 1) partial mediation (where the mediator accounts for some, but not all, of the effect of the independent variable on the dependent variable), and 2) complete mediation (where the effect of the independent variable on the dependent variable can be fully explained by the mediator).

Figure 4 presents the results of the mediation analysis with AD pathology as a mediator of the relationship between E/I ratios and cognitive performance and decline. Panel 4b illustrates whether the association between regional E/I ratios and global cognition at baseline is mediated by AD pathologies, while panel 4c explores the same relationship with cognitive decline as the outcome. In both cases, we did not observe significant mediator effects from NFT pathology. However, Aβ levels and Global AD did mediate the relationship between regional E/I ratios and cognitive scores. Significant mediation was found in brain regions such as the superior frontal gyrus, caudate, hippocampus, and putamen. Furthermore, we observed more cases of complete mediation for the slope of global cognition (cognitive decline) than for global cognition at baseline. Notably, Global AD values appeared to mediate the relationship more strongly than Aβ levels alone, further highlighting the importance of this combined metric in associations with cognitive decline.

Additionally, when we reversed the roles of biological variables, with E/I as a potential mediator of the association between AD pathologies and cognition, we found significant partial mediation (but not complete mediation) for global cognition in the left cerebellum crus II and left cerebellum VIIB, and for the slope of global cognition in the left superior parietal and left cerebellum VIIB only. It is important to note that reversing the mediation model (by swapping the independent and mediator variables) does not necessarily provide insight into the true direction of causality, as differences in model fit may reflect data variability rather than the underlying causal mechanism. However, the linear associations observed suggest a predominantly mediating role of AD pathologies, instead of E/I ratios, and underscore the importance of considering both pathologies and brain region-specific E/I in understanding cognitive outcomes.

### Molecular predictors of E/I imbalance highlight synaptic and immune dysfunction

Next, we identified transcriptomic and proteomic markers predictive of E/I ratios using Elastic-Net regression with a nested cross-validation framework (Figure 5a; see Statistical Analyses, *Molecular Predictors*). Only features quantified in both the SFG and ITG were included, with a covariate added to adjust for regional effects. Separate models were fit for transcriptomic and proteomic data, and overall model performance was quantified by the mean R² across 100 test folds (transcriptomics: R² = 0.31, proteomics: R² = 0.34). A final set of 89 genes and 101 proteins was obtained by prioritizing those features most consistently retained across folds, based on their selection frequency (Supplementary Table 6). The E/I ratio in both the SFG and ITG presented a positive relationship with most of the identified genes (85.4%) and proteins (67.3%). In these cases, higher gene expression or protein concentration correlated with a higher E/I ratio. Several genes and proteins consistently appeared across cross-validation folds, including *MET*, *LINC01122*, *SLC24A4*, *HMGXB4*, *GREB1L*, FMR1, PENK, B2M, CPNE9, RGS19, SLC9B2, and CADPS2_1 (Figure 5a). These molecules participate in a range of AD-related pathophysiological mechanims^70–76^, including synaptic dysfunction, neurodegeneration, chromatin remodeling, and neuroinflammation. Notably, several of identified molecular predictors such as *MET*, *SLC24A4,* FMR1, B2M, and CADPS2 have been directly linked to AD in previous studies^70–74^, supporting their association with the observed E/I imbalance.

**Figure 5.**
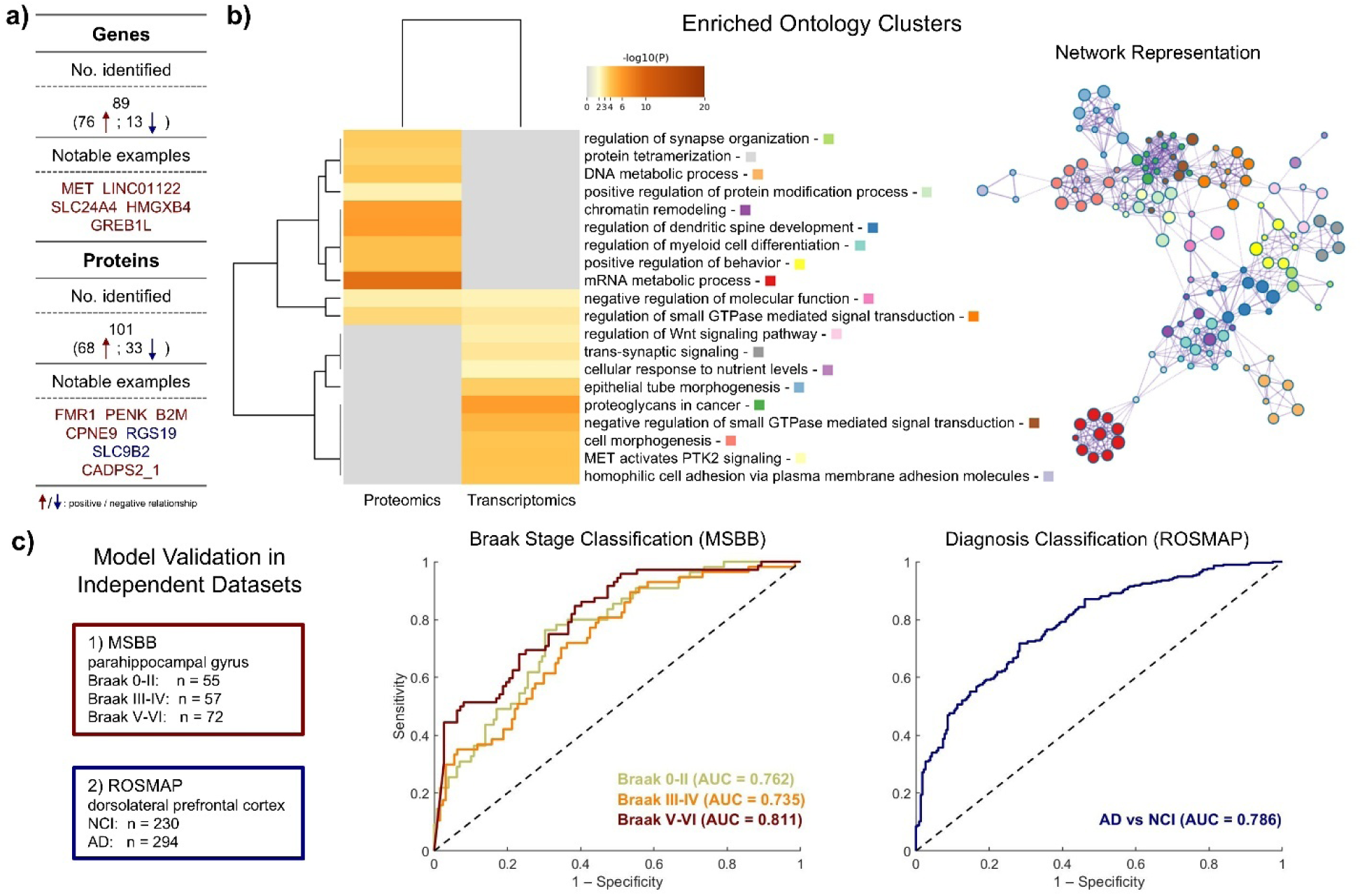
Molecular determinants of E/I ratios and enriched biological pathways. **a**) Summary of the transcriptomic and proteomic markers associated with E/I ratios in the superior frontal gyrus and inferior temporal gyrus, including the number of genes and proteins with positive and negative relationships to E/I imbalance, as well as the top predictors consistently identified across cross-validation folds. **b**) Pathway enrichment analysis of the identified predictors, displayed in a dendrogram. Heatmap cells are colored by their p-values, with white cells indicating no enrichment for that term in the corresponding list. The network visualization illustrates the interconnectedness of the molecular processes involved, with node size representing the number of genes in each pathway and edge thickness indicating pathway similarity. **c**) Receiver operating characteristic curves for classification performance in independent transcriptomic and proteomic datasets. Multiclass classification was performed on the Mount Sinai Brain Bank cohort using data from the parahippocampal gyrus to predict Braak stages (0–II, III–IV, V–VI). Data from the dorsolateral prefrontal cortex of an independent subset of ROSMAP participants were used to evaluate the predictive value of selected genes and proteins in distinguishing AD from NCI. Results shown correspond to a linear support vector machine classifier. These analyses assess the generalizability of selected molecular features across cohorts and brain regions. ROSMAP, Religious Orders Study and Memory and Aging Project; MSBB, Mount Sinai Brain Bank; AUC, area under receiver operating characteristic curve; AD, Alzheimer’s disease; NCI, no cognitive impairment.

Subsequently, we characterized the biological pathways enriched in the molecular predictor lists using Metascape (Supplementary Table 7). The results revealed several biological clusters, each represented by the term with the best p-value and displayed in a dendrogram in Figure 5b. The specific list of E/I ratio determinants (transcriptomics or proteomics) where each term is overrepresented is indicated in the dendogram. The top molecular processes are also visualized in a network layout, with node size proportional to the number of genes associated with each pathway and edge thickness reflecting the degree of similarity between pathways. This fully connected network emphasizes the tight relationship between the identified molecules and the biological processes in which they participate. According to their functional themes, the identified pathways can be categorized into the following groups: 1) Gene and Chromatin Regulation, including DNA and mRNA metabolic processes and chromatin remodeling; 2) Cell Morphology, Morphogenesis, and Differentiation, including cell morphogenesis, epithelial tube morphogenesis, homophilic cell adhesion via plasma membrane adhesion molecules, regulation of synapse organization, regulation of dendritic spine development, and regulation of myeloid cell differentiation; 3) Cell Signaling and Signal Transduction, including regulation of small GTPase-mediated signal transduction, regulation of the Wnt signaling pathway, trans-synaptic signaling, and cellular response to nutrient levels; and 4) Protein Modification and Metabolism, including positive regulation of protein modification processes, protein tetramerization, and negative regulation of molecular function. Previous studies have highlighted the individual relevance of many of these pathways in AD, particularly in the context of synaptic dysfunction, neuroinflammation, and cellular stress^76–87^. However, our comprehensive analysis underscores the interconnectedness of these basic cellular and molecular processes, which collectively contribute to AD pathogenesis.

### Molecular markers of E/I imbalance predict neuropathology and dementia across cohorts

To assess the generalizability of the identified network of genes and proteins predicting the E/I ratio, we performed external validation in two independent datasets comprising samples from distinct brain regions and cohorts (Figure 5c). Specifically, we used data from the Mount Sinai Brain Bank (MSBB) cohort^132^ (N = 184) with parahippocampal gyrus samples and a separate subset of ROSMAP participants (N = 524) with samples from the dorsolateral prefrontal cortex samples (DLPFC)^115,131^. In both datasets, all subjects with both gene and protein expression data were included. The outcomes were Braak stage (0–II, III–IV, V–VI) in MSBB and clinical diagnosis (NCI vs. AD) in ROSMAP. Molecular predictors were restricted to the set of genes and proteins overlapping with our previously identified features.

Several classification models were evaluated using 5-fold cross-validation, with the best performance achieved using the support vector machine (SVM) method. The multiclass Braak stage prediction (MSBB data) yielded robust areas under the receiver operating characteristic curve (AUC), with AUCs of 0.762 for Braak 0–II, 0.735 for Braak III–IV, and 0.811 for Braak V–VI. In the NCI vs AD classification (ROSMAP data), performance was similarly strong (AUC = 0.786). These externally validated results support that the identified transcriptomic and proteomic markers of E/I imbalance retain predictive value across different cohorts and brain regions, highlighting their relevance to AD pathophysiology and clinical deterioration.

### Impact of cell type distributions on the E/I ratio in AD

Finally, we assessed the relationship between the spatial E/I ratio map of AD—derived from regional comparisons of E/I ratios between AD subjects and NCI individuals—and the abundance of six major brain cell types estimated via gene expression data, as previously reported^88^. The cell types include neurons, astrocytes, oligodendrocytes, microglia, endothelial cells, and oligodendrocyte precursor cells (OPCs).

Interestingly, Spearman’s rank correlation revealed significant associations for microglia (r = 0.25) and OPCs (r = -0.37). Specifically, higher microglia abundance was linked to higher E/I ratios, while greater proportions of OPCs were associated with lower E/I ratios (Figure 6). Microglia contribute to synaptic clearance, pruning and neuroinflammation, which are critical processes underlying the balance between excitatory and inhibitory signaling in AD^89–91^. In addition, OPCs proliferate in face of myeline damage and are involved in maintenance and repairing of neural connections, potentially counteracting excitatory imbalances in the disease^92–94^. These findings support a role for microglia and OPCs in regional vulnerability to AD-associated neuronal dysfunction.

**Figure 6.**
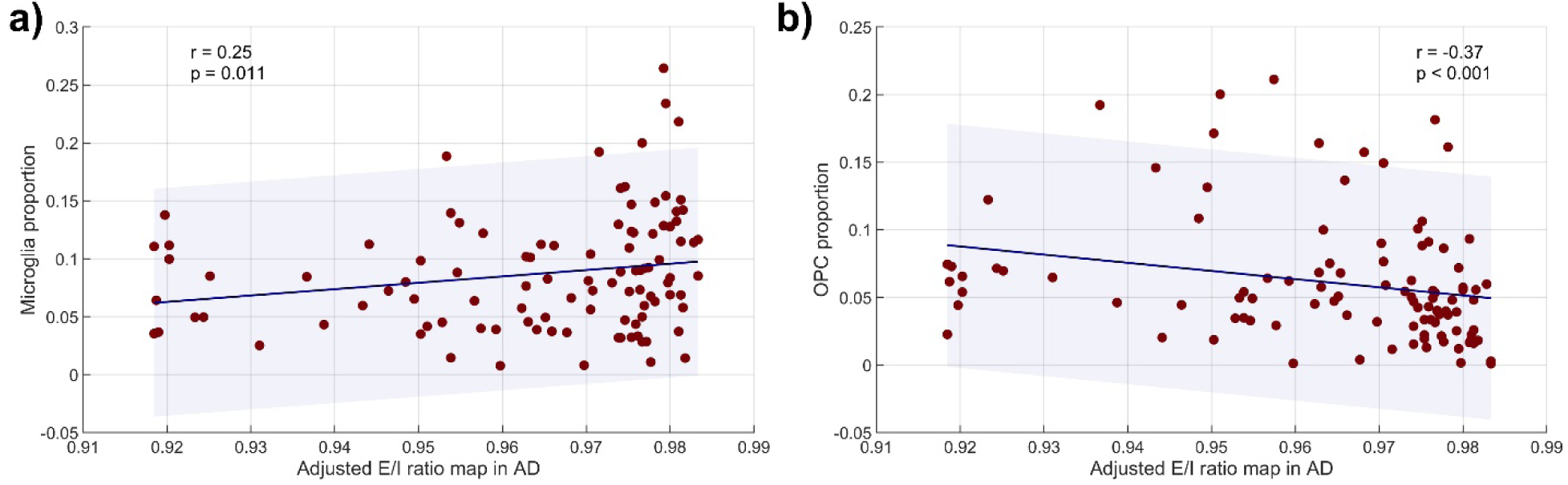
**Associations between across-brain cell abundance distributions and E/I ratios in AD**. Results of Spearman’s correlation analyses for the proportions of microglia (**a**) and OPCs (**b**). Error bands denote 95% confidence intervals. No significant relationships were found for neurons, astrocytes, oligodendrocytes, or endothelial cells.

## Discussion

Brain function emerges from complex interactions across multiple biological scales. However, the molecular mechanisms underlying brain dysfunction in individuals with AD dementia remain poorly understood. Two major challenges have hindered progress in this field: 1) the lack of large-scale datasets that simultaneously capture molecular profiles, brain activity, and clinical data from the same subjects, and 2) the difficulty in combining multi-scale data corresponding to diverse biological sources. Here, we developed a multi-scale computational framework that integrates ante-mortem resting-state fMRI data with post-mortem molecular omics from the same human participants. This approach allows us to interrogate the holistic relationship between neuronal E/I imbalance, cognitive decline and neuropathological features, and to identify molecular features associated with this imbalance.

Our findings suggest that regional E/I imbalance may significantly contribute to AD cognitive decline. Key AD-related regions such as the hippocampus, insula, lingual gyrus, caudate nucleus, and thalamus—known for their roles in memory, attention, and cognition^57–61,63^— emerged as important predictors of cognitive performance (Figure 2). The data reveals a complex, region-specific pattern though, with predominantly positive relationships between the estimated E/I ratios and cognitive scores. Regional E/I ratios may reflect the functional integrity of networks crucial for cognition, helping to determine a subject’s position on the continuum of cognitive resilience^95^. Interestingly, the relationship between regional E/I ratios in key regions, such as the left hippocampus, right lingual gyrus and right insula, and cognitive scores appears to be mediated by the global AD pathology burden (Figure 4).

The observed inverted U-shaped pattern of E/I ratios across stages of NFT pathology (Figure 3) reveals important dynamics in brain network dysfunction during AD progression. In the early stages (Braak 0-II), a lower E/I ratio may reflect the absence or onset of neurodegenerative processes^7^. As the disease progresses to the limbic stage (Braak III-IV), an increase in E/I balance may occur, driven by heightened excitatory signaling, decreased inhibition, or both, as the brain attempts to maintain cognitive function amidst growing neurodegeneration^2,20,102,103^. Interestingly, in a previous study integrating PET-measured brain distributions of Aβ and tau into personalized dynamic models, we identified the limbic stages as the critical phase in AD progression, where changes in electrophysiological activity first emerge^20^. This is also when cognitive decline becomes evident^64,65^. In the later stages (Braak V-VI), a new decrease in the E/I ratio could indicate widespread neuronal loss^2^, leading to a generalized reduction in neuronal activity and connectivity. The brain regions showing significant effects of Braak stage on the E/I ratio—such as the superior frontal gyrus, caudate, hippocampus, putamen, and thalamus—are particularly vulnerable to tau pathology and are also critical to higher-order cognitive functions^42,104–110^. These regions, heavily interconnected within key brain networks, may have their E/I balance more easily disrupted by the spread of tau pathology^2^. The lack of association with Aβ measures, such as the CERAD stages, points to tau’s disruptions to the microtubule network, synaptic transmission, and axonal transport as more influential than Aβ’s effects on extracellular processes^2^ in driving E/I imbalances. However, we also observed global (Aβ & tau) contributions in regions like the lingual gyrus (involved in visual processing and memory encoding^61^) and insula (associated with emotional and sensory processing^57^), which are part of Braak III-IV stages^42^. This global effect might also be linked to the increased excitation observed in the limbic phase of the disease. Notably, non-AD pathologies, often prevalent in aging populations, such as hippocampal sclerosis^43^ and atherosclerosis^45^ were also associated with significant alterations in E/I ratios, particularly in subcortical and temporo-frontal regions. These relationships suggest that disruptions to E/I balance are not exclusive to canonical AD pathology but may also emerge from comorbid processes that contribute to vascular compromise, neuroinflammation, or neuronal loss. Together, these findings point to a dynamic interplay between the pathological stage of the network and specific regional functional alterations.

In a subset of 87 individuals with rs-fMRI and both transcriptomic and proteomic data from the SFG and ITG, we identified several key genes and proteins whose expression levels were associated with regional E/I imbalances. Upregulation of more than 60% of the predictor genes/proteins was linked to higher E/I ratios. In the SFG, a significant increase in the E/I ratio was observed at Braak III-IV stages (Figure 3). For individuals at these stages of NFT involvement, it should be explored whether a therapeutic effect—by re-establishing healthy activity—could be achieved through precisely tuning the levels of genes/proteins associated with the E/I imbalance. Notably, the identified molecular features also demonstrated predictive utility for AD diagnosis and Braak staging in two independent cohorts with samples from different brain regions, reinforcing their broader relevance to AD-related pathophysiology (Figure 5c). While the specific roles of all the identified molecular determinants in AD require further investigation, several of the top contributors have already been directly linked to the disease. For example, genetic variations in *MET* and *SLC24A4* are associated with increased AD risk and may influence key neurological functions, including synaptic plasticity and ion homeostasis^70,71^. The proteins FMR1 and B2M are involved in the regulation of amyloid precursor protein (APP) and may co-aggregate in neuritic plaques^72,73^, a hallmark of AD pathology. Additionally, CADPS2, a calcium-dependent activator of secretion, plays a crucial role in synaptic vesicle dynamics; its dysfunction can impair neurotransmitter release and disrupt signaling^74,111^.

The enrichment analysis of the identified genes/proteins revealed significant associations with biological pathways thought to be critical for AD, particularly those governing synaptic signaling, cellular architecture, and immune response^7,19^ (Figure 5). Among the overrepresented biological processes was the regulation of Wnt signaling, which is essential for synaptic plasticity, neurogenesis, and blood-brain barrier integrity—with dysregulation implicated in AD^82^. Similarly, trans-synaptic signaling, a key process for neuronal communication, is impaired in AD, facilitating the spread of Aβ and tau^77^. Moreover, Aβ oligomers impair presynaptic organization by reducing the surface expression of β-neurexins (β-NRXs)^85^. Immune-related pathways, such as regulation of myeloid cell differentiation, were also enriched. This process may be important to microglial activation and the brain’s inflammatory response to AD pathology^81,112^. Additionally, small GTPases, like RhoA, are involved in microglial activation, and their dysregulation can contribute to chronic inflammation and neurodegeneration^87^. Finally, the regulation of mRNA metabolic processes was enriched in the group of molecules whose protein abundance was associated with E/I imbalances, with the lowest p-value among all identified biological processes. This finding suggests that not only gene transcription, but also the processing, stabilization, and translation of mRNA into proteins^78^, plays a crucial role in maintaining the balance of excitatory and inhibitory signaling. It may also point to potential therapeutic targets in AD for restoring proper protein synthesis pathways, particularly those involved in synaptic function and metabolic regulation. Given the connectedness of the identified molecular pathways, interventions targeting these processes could simultaneously benefit multiple biological domains.

Lastly, the association between the E/I ratio in AD and the abundance of microglia and oligodendrocyte precursor cells (OPCs) supports the idea that glial cells play critical roles in shaping regional neural activity in both health and disease. The cellular maps utilized in this study, derived from gene expression data in neurotypical brains^88^, provide a "by design" template for how microglia and OPCs distribute across regions. We found that regions with higher microglial abundance are associated with higher E/I ratios, while regions with greater OPC abundance tend to show lower E/I ratios in AD (Figure 6). Beyond their well-established role in myelination, OPCs are uniquely equipped to monitor neural activity and neurotransmitter release^93,94^. Recent evidence shows that OPCs can additionally engage in synaptic pruning, predominantly targeting excitatory synapses^92^, which may contribute to the lower E/I ratios observed in regions with high OPC abundance. In contrast, microglia, the brain’s resident macrophages^90,91^, also participate in synaptic pruning and may preferentially target inhibitory synapses in certain contexts^113,114^, potentially disrupting the E/I balance and leading to heightened excitation in AD. Moreover, microglial phenotypes may shift during disease progression, often driven by pathological stimuli, which could further modulate their impact on synaptic activity^89,90^. Although the precise mechanisms in AD remain unclear, microglia and OPCs may significantly contribute to the spatial vulnerability to AD-associated E/I imbalances in affected regions.

Despite the strengths of our approach, our conclusions are limited by a few methodological aspects. First, while our analysis included transcriptomic and proteomic data from the same individuals who underwent ante-mortem rs-fMRI, only two regions—the SFG and ITG—have been characterized to date in a considerable number of brains^31,115^. These region-specific insights provide valuable mechanistic understanding, but they fall short of offering a brain-wide view of the molecular processes that shape emerging brain activity patterns in AD. Future work that expands omics extraction to additional brain regions will be essential. Second, from a computational perspective, capturing the full complexity of interactions and spatial-temporal dynamics in living human brains remains a challenge^116^. Several refinements to the biophysical model could be implemented, such as incorporating subject-specific connectomes, adding more neuronal populations per brain region, and estimating inhibitory parameters^19,20^. Blood oxygenation level-dependent (BOLD) resting-state fMRI (rs-fMRI) signals are directly influenced by vascular and metabolic factors^117,118^, which were not individually estimated in this study. Additionally, models should incorporate astrocytic regulation of glutamatergic and GABAergic signaling to modulate excitatory and inhibitory synapses, respectively, which is crucial for maintaining proper E/I balance^119,120^. Nevertheless, more detailed models typically increase the computational costs of simulating neuronal activity. The computational framework used in this study significantly accelerated fMRI signal simulations by leveraging machine learning techniques, which can now run on a standard desktop computer (whereas previously, simulating this complex dynamical system with many differential equations required substantial memory and time on a computing cluster^20^). This improvement should enable the identification of additional parameters in higher-dimensional systems in future studies.

Our study contributes to the growing effort to understand the molecular determinants of macroscopic brain activity, including functional connectivity and inter-individual structural variations^31^. By integrating multi-omics data with resting-state fMRI, we offer a more nuanced view of how brain architecture, AD pathology, and molecular disruptions across key processes shape regional E/I imbalances, ultimately influencing cognitive decline or resilience. Overall, our findings provide integrative molecular insights into the biological mechanisms underlying brain function alterations in AD, revealing potential targets for therapeutic intervention.

## Methods

### ROSMAP cohort

#### Ethics statement

The Religious Orders Study (ROS) and the Rush Memory and Aging Project Study (MAP)^32,33^ were each approved by an Institutional Review Board (IRB) of Rush University Medical Center. All participants enrolled without known dementia and signed an informed consent and Anatomical Gift Act agreeing to annual detailed clinical evaluation and post-mortem brain donation. In addition, they signed a repository consent allowing their data to be shared. Data documentation and sharing documents can be obtained at https://www.radc.rush.edu. The study was conducted according to Good Clinical Practice guidelines, the Declaration of Helsinki, and IRBs. Study subjects and/or authorized representatives gave written informed consent at the time of enrollment for sample collection and completed questionnaires approved by each participating site IRB.

#### Participants

The study involved 156 subjects from ROSMAP, all of whom underwent ante-mortem structural MRI and resting-state fMRI, as well as post-mortem neuropathological evaluations. The latest available magnetic resonance images, closest to the time of death, were utilized. Among these subjects with available neuroimaging, 87 had both gene expression and protein abundance data generated from samples in the superior frontal gyrus (SFG) and inferior temporal gyrus (ITG).

#### Clinical diagnoses

Each participant underwent a battery of 21 cognitive performance tests each year^121^. Of these, 17 tests were used to generate level of and change in global cognition and five cognitive domains: episodic, semantic, and working memory, perceptual speed, and visuospatial ability^122,123^. The tests also informed on clinical diagnoses at the time of death including dementia and Alzheimer’s dementia (AD-dementia), mild cognitive impairment (MCI), or no cognitive impairment (NCI) based on their clinical status^34–36^.

#### Neuroimaging

Structural scans were obtained using T1-weighted MRI across various scanners, including a 1.5T GE system and 3T Siemens and Philips systems, all with a resolution of 1 mm³. Detailed scanner protocols are available online (https://www.radc.rush.edu/docs/var/scannerProtocols.htm) and have been published elsewhere^31^. Preprocessing steps included non-uniformity correction, skull stripping, spatial normalization to the MNI152 template, and tissue segmentation, all performed with advanced normalization tools^31^. To construct individual gray matter parcellations, each individual’s native space gray matter was divided into 70 cortical regions, 16 subcortical nuclei, and 18 cerebellar lobules, based on the Desikan–Killiany Atlas and AAL cerebellar lobules^124^.

Resting state fMRIs were acquired on multiple scanners, including a 1.5T GE Signa scanner (70 mm³ voxel size, TR/TE = 2000/33 ms, flip angle 85°) and a 3T Siemens Magnetom Trio (36 mm³ voxel size, TR/TE = 3000/30 ms, flip angle 80°). Detailed scanner protocols are available online (https://www.radc.rush.edu/docs/var/scannerProtocols.htm) and have been published elsewhere^31^. The preprocessing followed a robust pipeline^31^, starting with CuBIDS to identify scans with deviant parameters and reduce scan heterogeneity. The data were then processed with fMRIPrep (v.20.2.3) for realignment, slice-time correction, and distortion correction (if applicable). fMRI volumes were co-registered to T1-weighted images using FLIRT. Confound regression was performed using the eXtensible Connectivity Pipeline (XCP; v.0.0.4), including bandpass filtering (0.01–0.08 Hz), despiking, and regression of 36 motion attributes, physiological parameters and a global signal. We then computed the amplitude of low-frequency fluctuations (ALFF, 0.01–0.08 Hz) and averaged them over all voxels belonging to a brain region to yield a single value per region^51^. ALFF serves as an indicator of spontaneous neuronal activity^48,49^.

#### Neuropathological assessments

All participants underwent neuropathological evaluations, as previously described^32,41,42^. Braak staging was used as a semiquantitative measure of neurofibrillary tangle (NFT) severity. Stages I and II indicate NFTs confined primarily to the entorhinal cortex, stages III and IV suggest involvement of the limbic regions (e.g., hippocampus), and stages V and VI reflect moderate to severe neocortical involvement. Neuritic plaque density was assessed using the Consortium to Establish a Registry for Alzheimer’s Disease (CERAD) criteria, with a semiquantitative scoring system for plaque burden in neocortical regions. AD diagnoses were based on the CERAD score, with classifications ranging from no AD to possible, probable, or definite AD (defined by frequent neuritic plaques in one or more neocortical regions). Both Braak staging and CERAD scoring were guided by a neuropathologist’s opinion and an established algorithm^32,41,42^. Additionally, regional counts of neuritic plaques, diffuse plaques, and NFTs were obtained from microscopic examination of silver-stained slides across five regions: midfrontal cortex, midtemporal cortex, inferior parietal cortex, entorhinal cortex, and hippocampus. Scaled values for each pathology were averaged to compute a global quantitative summary of AD pathology (gpath)^39,40^.

Common age-related pathologies were categorized into stages, as reported in previous studies^43–47^. The assessments included Lewy body disease (not present, nigral-predominant, limbic-type, neocortical-type), TDP-43 pathology (none, amygdala, amygdala + limbic, amygdala + limbic + neocortical), hippocampal sclerosis (not present, present), arteriolosclerosis (none, mild, moderate, severe), cerebral atherosclerosis (none, mild, moderate, severe) and presence of chronic macroinfarctions (none, one or more) and microinfarctions (none, one or more).

#### Proteomics

Multiplex tandem mass tag mass spectrometry (TMT-MS) was used to analyze 7,788 proteins from brain tissue samples (100 mg each) collected from 87 ROSMAP participants, specifically from the SFG and ITG. Following homogenization and protein quantification, proteins were digested and labeled with TMT using an 11-plex kit. High-pH offline fractionation yielded 24 pooled fractions, which were analyzed via liquid chromatography-tandem mass spectrometry on an Orbitrap Fusion mass spectrometer. Data processing with Proteome Discoverer identified 10,426 high-confidence proteins, with final analyses including only proteins quantified in over 50% of samples. Batch correction and outlier detection were performed to ensure data integrity. Detailed steps can be found in a previous publication^115^.

#### Gene Expression

RNA was extracted from brain tissue samples collected from the SFG and ITG using the Chemagic RNA tissue kit. Quality and concentration were assessed with the Fragment Analyzer and Qubit assay. For RNA-Seq, 500 ng of total RNA underwent rRNA depletion and library preparation using TruSeq stranded libraries. The libraries, averaging 330-370 bp, were sequenced on a NovaSeq 6000, generating 80-100 million paired-end reads (2x150 bp). RNA-Seq data processing involved quality control (QC), gene and transcript quantification, and 3’ UTR quantification. QC included alignment with STAR to the human reference genome (Gencode Release 27 GRCh38), while raw counts were generated using Kallisto. Data were normalized as Counts Per Million (CPM) and adjusted for confounding factors using voom/limma. Only genes with mean log2(CPM) > 2 were retained. Detailed steps can be found in a previous publication^115^.

### Reference data

#### Anatomical connectivity

The connectivity matrix was obtained using DSI Studio (http://dsi-studio.labsolver.org). A group average template was constructed from a total of 1065 subjects in the Human Connectome Project^125^. We employed a multishell diffusion scheme with b-values of 990, 1985, and 2980 s/mm². The number of diffusion sampling directions was 90 for each shell. The in-plane resolution and slice thickness were both set to 1.25 mm. An automatic quality control routine verified the b-table for accuracy^126^. The diffusion data were reconstructed in the MNI space using q-space diffeomorphic reconstruction to obtain the spin distribution function^127^. A diffusion sampling length ratio of 1.7 was used. The output resolution is 1 mm isotropic. Restricted diffusion was quantified using restricted diffusion imaging^128^. A deterministic fiber tracking algorithm^129^ was employed, enhanced by augmented tracking strategies^130^ to improve reproducibility. The anisotropy threshold was randomly selected between 0.5 and 0.7, while the angular threshold ranged from 45 to 90 degrees. The step size matched the voxel spacing. Tracks shorter than 30 mm or longer than 200 mm were discarded, with a total of 1,000,000 seeds placed. Shape analysis^130^ was performed to derive shape metrics for tractography. The connectivity matrix was calculated based on the count of connecting pathways in the selected brain parcellation.

### Cell abundance maps

Cell type maps were previously generated^88^ by estimating the proportions of six major cell types (neurons, astrocytes, oligodendrocytes, microglia, endothelial cells, and oligodendrocyte precursor cells [OPCs]) in gray matter (GM). This was done using the AHBA transcriptome atlas, which provides gene expression data from multiple brain regions across six neurotypical human donors^26^. Genetic markers for each cell type were selected from the BRETIGEA human brain marker gene set and matched to the AHBA gene probes. Cell type proportions were then estimated using eigengene decomposition, normalized by GM density, and registered into MNI space for analysis. The resulting cell densities were averaged over 104 anatomical regions defined by the Desikan–Killiany cortical atlas, as well as subcortical regions and AAL cerebellar lobules. Detailed steps can be found in a previous publication^88^ (https://github.com/neuropm-lab/cellmaps).

### Independent validation data

#### ROSMAP (different subjects and samples)

Additional postmortem brain transcriptomic and proteomic data from the dorsolateral prefrontal cortex of ROSMAP participants were obtained from the Accelerating Medicines Partnership Alzheimer’s Disease (AMP-AD) Knowledge Portal (www.synapse.org). Bulk RNA-seq data (Synapse ID: syn3388564) were available for 17,309 genes, generated using next-generation sequencing (NGS), with gene counts estimated via Kallisto and normalized as previously described^115^. Proteomic data (Synapse ID: syn17015098) included quantification of 10,030 proteins using tandem mass tag (TMT) mass spectrometry, as detailed in prior work^131^. Of all available subjects, 38 were included in the main cohort of this study and were subsequently excluded from the validation analysis, leaving 524 AD and NCI independent subjects for analysis.

#### MSBB

Postmortem transcriptomic (RNA-seq) and proteomic (TMT) data from the parahippocampal gyrus of Mount Sinai Brain Bank (MSBB) participants were also obtained (syn3159438). Preprocessing followed established protocols comparable to those used in ROSMAP and has been described previously^132^. From this cohort, 184 subjects with both data types and neuropathological information were considered.

### Estimating neuronal activity alterations

We combined a Wilson-Cowan (WC) excitatory and inhibitory network model with metabolic and hemodynamic transformations to simulate brain dynamics^19,20^. The WC model represents each brain region’s activity as a function of excitatory and inhibitory interactions, where the firing threshold (θ) controls neuronal excitability within each region^55,133,134^. The global coupling strength (η) governs communication between regions, determining how activity propagates across the brain^52,135^. Following established methods, we computed the excitatory/inhibitory (E/I) ratio for the excitatory populations as the time average of the relative contributions of excitatory and inhibitory synaptic input currents to sustained oscillations^1,52–54,136^. Detailed formulations of these calculations can be found in previous publications^19,20^ and the Supplementary Text.

To estimate η and θ for each subject, we simulate brain activity^25,137^ and compare the resulting BOLD signals to actual data. The goal is to minimize the difference between simulated and real BOLD signals, ensuring that the model accurately reflects individual-specific brain dynamics^19,20^. We implemented a pipeline utilizing grid simulations, pre-trained feedforward neural network architectures and surrogate optimization techniques (Supplementary Figure 1).

#### Grid simulations of the biophysical model

Simulations were conducted using a previously-published^138,139^ MATLAB (The MathWorks Inc., Natick, MA, USA) function designed to model neural mass dynamics and BOLD signal responses based on specified parameters (see also Supplementary Text-Table 1). A fourth-order Runge-Kutta method (ode45) was used to numerically solve the differential equations governing the system dynamics. Simulations were run across a grid defined by the firing threshold and coupling strength parameters, with transient periods accounted for to stabilize initial conditions. BOLD signals were calculated from neural activities, and the amplitudes of low-frequency fluctuations (ALFF) were extracted for each parameter configuration.

#### Accelerated simulations assisted by machine learning

We utilized a feedforward neural network (FNN) implemented in PyTorch^140^ for predictive modeling of the ALFF values across combinations of firing threshold (θ) and coupling strength (η). Two separate FNN models were used, one for each fMRI acquisition protocol, as the generated biophysical signals are influenced by the specific settings of each protocol. The biophysical simulations on the parameter grid were divided into training, validation, and test sets using a stratified random split to ensure representative distributions. The model architectures included multiple hidden layers with varying configurations, such as dropout rates, learning rates, batch sizes, weight decay, and the number of epochs, which were systematically explored through hyperparameter tuning. Training was carried out using the Adam optimizer with a mean squared error loss function. We monitored training and validation losses to evaluate model performance and prevent overfitting or underfitting. After training, the best-performing configurations were identified based on validation loss, and the corresponding model weights were saved for further evaluation on the test set (Supplementary Figures 3 and 4). The optimal model configurations had the following parameters: five hidden layers with sizes 1024, 512, 256, 128, and 64 neurons; a learning rate of 0.001; no dropout; weight decay of 0; a batch size of 32; 400 training epochs for the 1.5 T MRI protocol and 600 training epochs for the 3 T MRI protocol. The trained FNN facilitates fast and reliable evaluations of the computationally intensive biophysical model.

#### Parameter Identification

The gp_minimize function from the skopt library^141^ in Python was used for parameter optimization. The loss function minimized the correlation difference between simulated and subject-specific ALFF values^20^. For each iteration, we evaluated the ALFF distribution for the current firing threshold and coupling strength using the selected FNN architecture and weights. Parameter bounds were defined based on preliminary evaluations of BOLD signals generated with the biophysical model (Supplementary Figures 3 and 4). For all subjects, we conducted 200 surrogate optimization iterations, saving the parameter sets (θ, η) with the lowest objective function values to use in new simulations that reconstruct the E/I ratio brain distributions for each subject.

### Statistical analyses

#### Prediction of cognitive scores

We analyzed cognitive scores, including global cognition, five cognitive domains, and their longitudinal slopes, in relation to regional E/I ratio values. Elastic net regression^56^ (MATLAB’s *lasso.m* function, with regularization mixing parameter α = 0.5) was used to predict each score, with covariates sex, age, years of education, and MRI protocol type residualized from both predictors and outcomes. For each outcome, the model was fit using 10-fold cross-validation to select the optimal regularization strength (λ). To assess the statistical significance of regional coefficients, we ran 1,000 permutations per outcome score, randomly shuffling the residualized outcome and refitting the model using the λ selected from the original fit. This yielded null distributions of coefficients per region, from which empirical p-values were computed. Multiple comparisons were corrected using the max-T method to control the family-wise error rate (FWER).

#### Pathological effects

We assessed the association between neuropathological measures (amyloid, tau, global pathology) and regional E/I ratios using partial correlation analysis (*partialcorr.m*) controlling for age, sex, years of education, and MRI protocol type. To correct for multiple comparisons across the 104 brain regions, we applied a permutation-based max-T procedure (1,000 permutations). For each permutation, the neuropathological variable was shuffled, partial correlations were recomputed across all regions, and the maximum Fisher z-transformed correlation was stored. FWER-corrected p-values were computed as the proportion of permutations in which the permuted maximum statistic exceeded the observed Fisher Z-value, with significance defined at 0.05.

We evaluated whether regional E/I ratios differ significantly by levels of neuropathological burden using a multi-way analysis of variance (*anovan.m*), including both categorical and continuous predictors. Independent variables included semi-quantitative staging measures (CERAD scores, Braak stages), non-AD-specific pathologies (Lewy bodies, TDP-43, hippocampal sclerosis, arteriolosclerosis, cerebral amyloid angiopathy, atherosclerosis, infarcts), and demographic covariates (age, sex, education, MRI protocol). To assess effect sizes, we computed η² values for each predictor and region. To correct for multiple comparisons, we implemented a permutation-based max-F procedure using 1,000 permutations, in which each predictor was independently shuffled and models refit. FWER-corrected p-values were calculated as the proportion of maximum permuted F-statistics exceeding the observed F-statistic for each variable and region, with 0.05 used as the significance threshold.

#### Mediation analysis

We investigated the relationship between regional E/I ratios, AD pathologies, and global cognitive scores using the Mediation Toolbox (https://github.com/canlab/MediationToolbox)^67–69^. We performed 10,000 bootstrap samples to estimate the indirect effect of the mediator in the relationship between the independent and dependent variables, allowing us to estimate confidence intervals and p-values for the mediation effect. Additionally, we employed the toolbox’s robust regression option to account for potential outliers in the data. The analysis was adjusted for covariates including age, sex, education years, and MRI protocol.

#### Molecular predictors

We implemented Elastic-Net regression^56^ (*lasso.m*) to identify transcriptomic and proteomic markers predictive of E/I ratios in the SFG and ITG. The model balanced L1 and L2 penalties (*lasso.m*, α = 0.5) and optimized the regularization parameter λ over a logarithmic grid from 10⁻³ to 10¹. Only genes and proteins quantified in both brain regions were included. For each omic type (transcriptomics and proteomics), data from SFG and ITG were modeled simultaneously with a ’region’ covariate to adjust for regional effects. A nested cross-validation scheme was used to assess model performance and feature stability^142^: the outer loop consisted of 100-fold Monte Carlo CV with random splits, reserving 20% of data as independent test sets while maintaining patient stratification to avoid leakage. The inner loop performed 5-fold CV within training data to select the optimal λ minimizing mean squared error. For each outer fold, the model was trained on training data, selecting features with non-zero coefficients at the optimal lambda. Predictions on held-out test sets were used to calculate R², which was averaged across the 100 outer folds to quantify model performance. Selected features for each omic type were aggregated across all folds, with those in the top 25th percentile by selection frequency retained for downstream multi-omic integration and pathway analyses (Supplementary Table 6). This threshold prioritizes consistently selected markers and reduces the risk of missing key biological processes by retaining too few molecular candidates or introducing noise by retaining too many^143^.

We performed functional enrichment analyses using Metascape^143^, a web-based platform that integrates multiple biological databases, including KEGG, GO, Reactome, and others (accessed October 30, 2024). Metascape identifies significantly enriched ontology terms in gene lists through hypergeometric tests, applying the Benjamini-Hochberg correction for multiple comparisons (q < 0.05). To minimize redundancy, Kappa-statistical similarities among enriched terms were calculated, followed by hierarchical clustering with a 0.3 threshold to define clusters. Enrichment results were visualized in a network format, where terms are represented as nodes with sizes proportional to the number of genes they contain and colors indicating cluster identity. Terms with a similarity score > 0.3 were connected by edges, with edge thickness reflecting similarity strength.

We also evaluated the predictive value of the identified E/I-related gene and protein features in independent datasets: an unseen subset of ROSMAP participants with DLPFC transcriptomic and proteomic data, and the MSBB cohort with matched data from the parahippocampal gyrus (PHG). In the ROSMAP-DLPFC dataset, the classification task was binary (AD vs NCI clinical diagnosis), using 78 genes and 93 proteins whose expression levels were quantified in both this dataset and the original cohort, with adjustment for age, sex, and years of education. In the MSBB-PHG cohort, a multiclass classification task was defined to predict Braak stages (0–II, III–IV, V–VI), using 86 genes and 90 proteins. Multiple supervised machine learning models were trained and evaluated, including support vector machines (SVMs), k-nearest neighbors (k-NN), decision trees, ensemble methods, and neural networks. Hyperparameter optimization was performed using internal cross-validation. Model performance was evaluated using 5-fold cross-validation, and classification accuracy was used as the primary criterion for model selection. Models included feature standardization prior to training. A Gaussian kernel SVM yielded the highest accuracy in the ROSMAP-DLPFC dataset, while a linear SVM was selected as the best-performing model in the MSBB-PHG dataset. Receiver operating characteristic (ROC) curves were generated from the cross-validation results to assess classification performance.

#### Cellular correlates

We used Spearman’s rank correlation (*corr.m*) to examine the relationship between the distribution of various cell types (neurons, astrocytes, oligodendrocytes, microglia, endothelial cells, and OPCs) and an AD-specific E/I ratio map across brain regions. The AD map was generated by applying rank sum tests to compare the E/I ratio between the AD-dementia and NCI diagnostic groups in each region, adjusting for age, sex, and MRI protocol.

## Data and code availability

All the materials needed to evaluate the conclusions in the study are provided in the article, the Supplementary Materials and/or referenced publications. The analyses used standard Python and MATLAB libraries, which have been described^138,139^. Detailed ROSMAP neuropathological, clinical, neuroimaging and gene expression data are available at the RADC Research Resource Sharing Hub (radc.rush.edu), pending scientific review and a completed material transfer agreement (see radc.rush.edu/requests.htm). Protein data are available through https://adknowl-edgeportal.synapse.org/.

## Acknowledgements

We are grateful to the participants in the Religious Order Study (ROS), and the Rush Memory and Aging Project (MAP) cohorts. LSR was partially supported by funding from the Fonds de recherche du Québec – Santé. This project was partly supported by the following awards to YIM: the Canada Research Chair tier-2, the CIHR Project Grant 2020, and the Transformational Research in AD 2020. In addition, we used the computational infrastructure of the McConnell Brain Imaging Center at the Montreal Neurological Institute, supported in part by the Brain Canada Foundation, through the Canada Brain Research Fund, with the financial support of Health Canada and sponsors. ROSMAP data was obtained thanks to funding by the National Institutes of Health to DAB (P30AG10161, P30AG72975, R01AG15819, R01AG17917, U01AG46152 and U01AG61356), and the Paul M Angell Family Foundation. The results published here are in whole or in part based on data obtained from the AD Knowledge Portal (https://adknowledgeportal.org/). The Mount Sinai Brain Bank (MSBB) transcriptomic and proteomic data were generated from postmortem brain tissue collected at the Mount Sinai VA Medical Center Brain Bank, provided by Dr. Eric Schadt. MSBB Proteomic data were generated by Dr. Allan Levey from Emory University.

