## Supplementary Information for "Molecular and neuropathological determinants of neuronal dysfunction in Alzheimer’s disease"

**Supplementary Table 1.** Demographics of the sample

| | NCI | MCI | p | AD | p |
|---|---|---|---|---|---|
| Number of individuals (% of total) | 69 (44.2) | 27 (17.3) | - | 60 (38.5) | - |
| Age (yrs), mean (s.d.) | 81.01 (6.17) | 83.02 (6.34) | 0.167 | 82.09 (4.59) | 0.259 |
| Female, N (%) | 51 (73.9) | 20 (74.1) | 1.000 | 47 (78.3) | 0.680 |
| Education (yrs), mean (s.d.) | 15.30 (3.06) | 15.22 (2.47) | 0.892 | 15.55 (2.54) | 0.619 |
| *APOE* ε4 carriers, N (%) | 10 (14.5) | 1 (3.7) | 0.168 | 18 (30.0) | 0.054 |
| MMSE, mean (s.d.) | 27.82 (1.53) | 25.03 (3.55) | < 0.001 | 13.88 (8.71) | < 0.001 |
| CERAD “yes”, N (%) | 33 (47.8) | 21 (77.8) | 0.011 | 56 (93.3) | < 0.001 |
| Braak ≥ 3, N (%) | 52 (75.4) | 25 (92.6) | 0.086 | 56 (93.3) | 0.007 |

The reported p-values are for comparisons to cognitively no cognitive impairment (NCI) subjects. P-values for age, education and MMSE indicate values assessed with two-sided independent-samples t-tests. For the resting variables (sex, *APOE* ε4 status, CERAD and Braak), Fischer exact tests were performed. NCI, no cognitive impairment; MCI, mild cognitive impairment; AD, Alzheimer’s disease-dementia; *APOE* ε4, apolipoprotein epsilon 4; MMSE, Mini-Mental State examination; CERAD “yes”, moderate or frequent neuritic plaques in one or more neocortical regions; Braak ≥ 3, limbic or neocortical neurofibrillary tangle pathology.

**Supplementary Table 2.** Demographics of the subsample with gene expression and protein data

| | NCI | MCI | p | AD | p |
|---|---|---|---|---|---|
| Number of individuals (% of total) | 47 (54.0) | 13 (14.9) | - | 27 (31.1) | - |
| Age (yrs), mean (s.d.) | 80.95 (6.36) | 85.01 (6.92) | 0.073 | 81.7 (4.81) | 0.566 |
| Female, N (%) | 34 (72.3) | 12 (92.3) | 0.264 | 21 (77.8) | 0.783 |
| Education (yrs), mean (s.d.) | 15.23 (3.32) | 15.31 (1.89) | 0.918 | 15.41 (2.59) | 0.804 |
| *APOE* ε4 carriers, N (%) | 5 (10.6) | 1 (7.7) | 1.000 | 9 (33.3) | 0.029 |
| MMSE, mean (s.d.) | 27.78 (1.57) | 25.71 (2.77) | 0.021 | 13.42 (8.72) | < 0.001 |
| CERAD "yes", N (%) | 21 (44.7) | 11 (84.6) | 0.013 | 25 (92.6) | < 0.001 |
| Braak ≥ 3, N (%) | 38 (80.9) | 13 (100.0) | 0.183 | 26 (96.3) | 0.082 |

The reported p-values are for comparisons to cognitively no cognitive impairment (NCI) subjects. P-values for age, education and MMSE indicate values assessed with two-sided independent-samples t-tests. For the resting variables (sex, *APOE* ε4 status, CERAD and Braak), Fischer exact tests were performed. NCI, no cognitive impairment; MCI, mild cognitive impairment; AD, Alzheimer's disease-dementia; *APOE* ε4, apolipoprotein epsilon 4; MMSE, Mini-Mental State examination; CERAD "yes", moderate or frequent neuritic plaques in one or more neocortical regions; Braak ≥ 3, limbic or neocortical neurofibrillary tangle pathology.

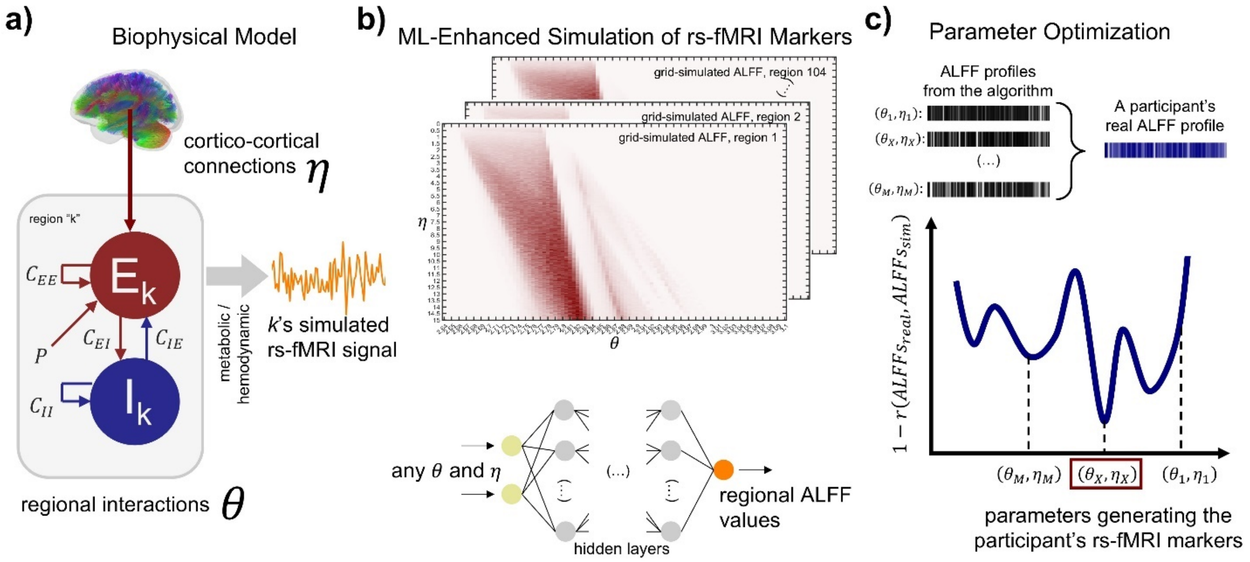


**Supplementary Figure 1. Modeling, simulating, and optimizing brain dynamics using a Wilson-Cowan excitatory-inhibitory network and machine learning. a**) Schematic representation of the WC network model for each brain region, where neuronal activity is governed by excitatory (E) and inhibitory (I) interactions. Brain dynamics are influenced by two key parameters: the firing threshold (θ), which controls neuronal excitability, and the global coupling strength (η), which regulates inter-regional communication. The resulting BOLD signals depend on these parameters. **b**) Overview of the simulation pipeline. Top: Grid simulations of the biophysical model were performed across a parameter space defined by θ and η, generating regional ALFF values for different combinations. Bottom: Accelerated simulations using a feedforward neural network (FNN) trained on the grid simulation results. The neural network takes θ and η as input and outputs predicted regional ALFF values, facilitating fast and accurate evaluations of the computationally expensive biophysical model. **c**) Parameter identification using surrogate optimization. ALFF values for various θ and η combinations are shown, along with the correlation distance between simulated and real ALFF distributions for each subject. The optimal parameter set, corresponding to the lowest correlation distance, is retained as the most likely set of parameters that generate the subject's real BOLD signal.

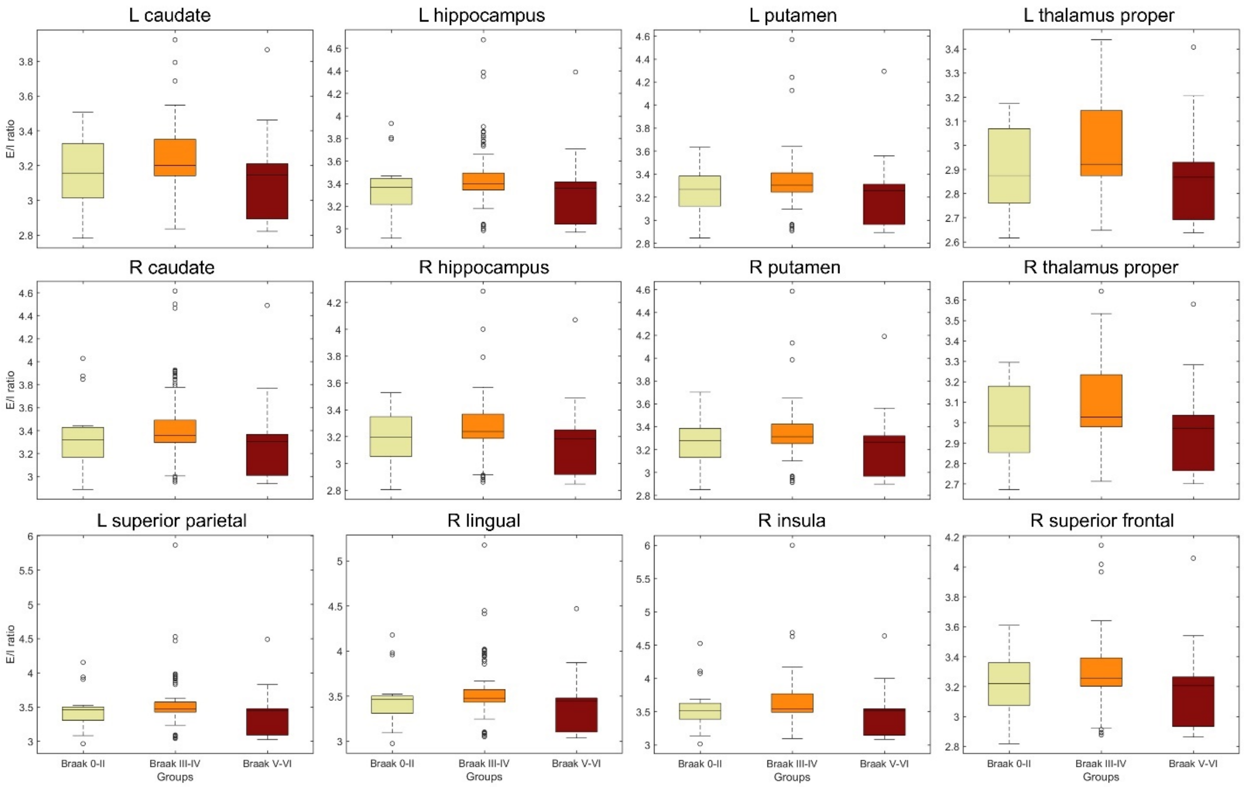


**Supplementary Figure 2. E/I ratio by NFT severity across regions with significant relationships.** Box-and-whisker plot illustrating the inverted U-shaped pattern of E/I ratios across Braak stages for all regions where significant relationships with Braak stage were identified. In the box-and-whisker plot, the central line indicates the group median, with the bottom and top edges of each box representing the 25th and 75th percentiles, respectively. Whiskers extend to the maximum and minimum values, while outliers are plotted individually as circles. The pattern is consistent across these regions, showing lower E/I ratios at Braak stages 0-II, higher ratios at Braak stages III-IV (limbic), and decreasing again at Braak stages V-VI (neocortical).

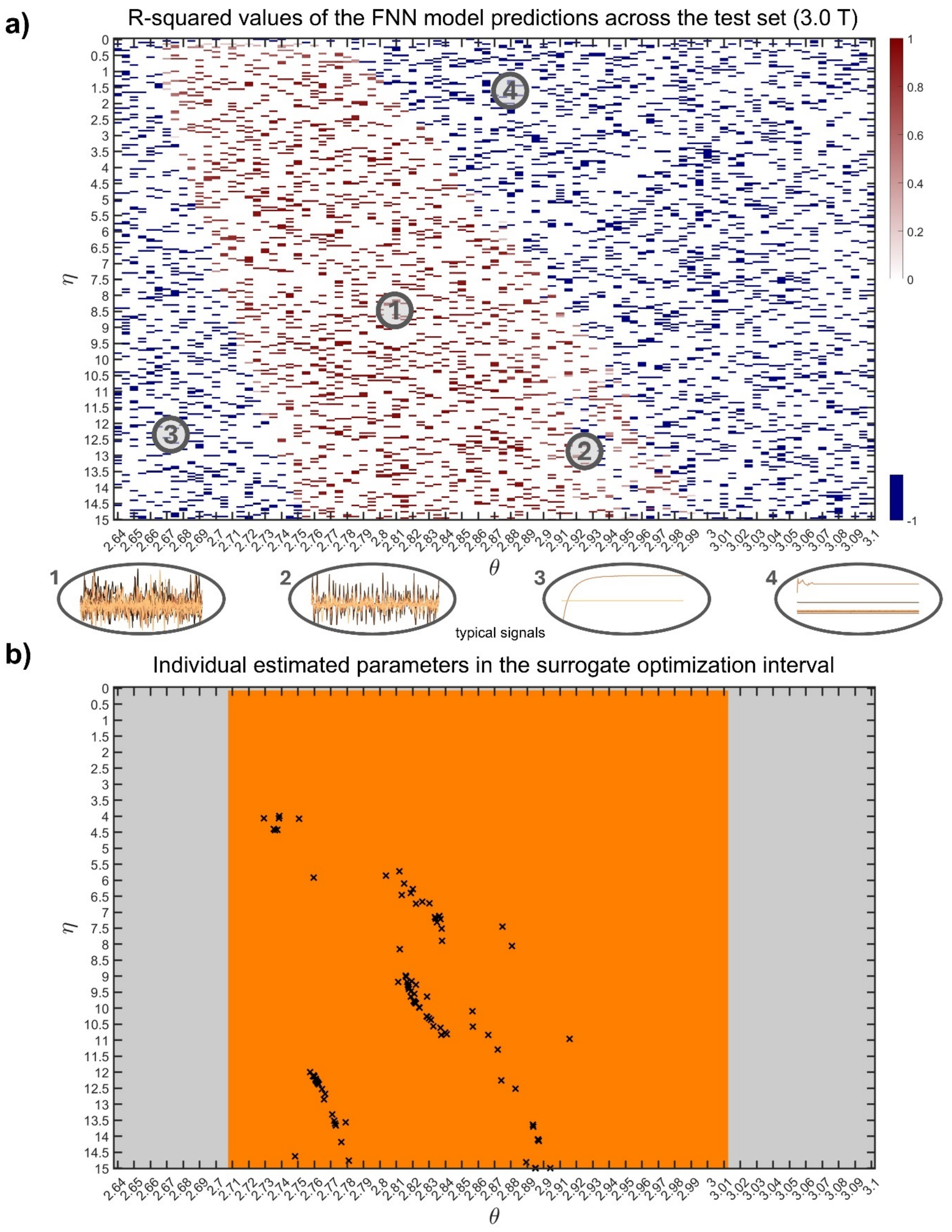


**Supplementary Figure 3. FNN training and parameter optimization results for the 3T MRI protocol. a**) R-squared ($R^2$) values of the FNN model's predictions across the test set, plotted on a grid of coupling strength (η) and firing threshold (θ). The $R^2$ values are shown for predictions based on the FNN, compared to simulated ALFF values across different η-θ combinations. Four representative points are highlighted, showing the fMRI signals for each set of η and θ. High $R^2$ values are observed in regions where the model successfully predicts fMRI signals that resemble real ones, while low $R^2$ values are associated with parameter sets that fail to produce realistic signals, such as the absence of oscillations (all negative $R^2$ values, denoting poor fit, were set to -1 in the figure). **b**) Estimated parameters (marked as crosses) for each subject, obtained through surrogate optimization, are plotted on the same η-θ grid. An orange box highlights the region of the parameter space explored during optimization. This region corresponds to the surrogate optimization interval, within which the parameter sets are considered to best match the subject-specific fMRI data for the 3T protocol.

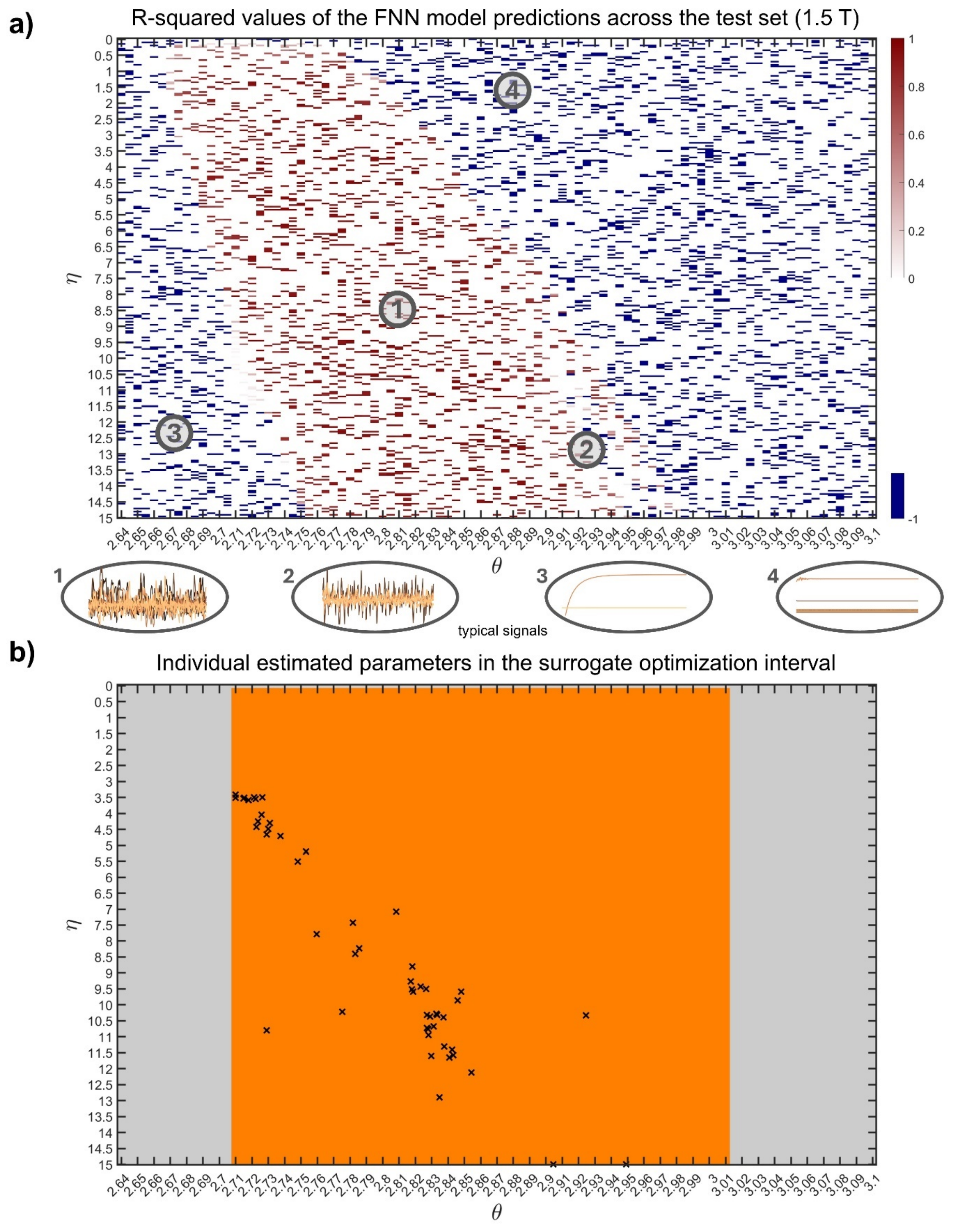


**Supplementary Figure 4. FNN training and parameter optimization results for the 1.5T MRI protocol. a**) R-squared ($R^2$) values of the FNN model's predictions across the test set, plotted on a grid of coupling strength (η) and firing threshold (θ). The $R^2$ values are shown for predictions based on the FNN, compared to simulated ALFF values across different η-θ combinations. Four representative points are highlighted, showing the fMRI signals for each set of $\eta$ and $\theta$. High $R^2$ values are observed in regions where the model successfully predicts fMRI signals that resemble real ones, while low $R^2$ values are associated with parameter sets that fail to produce realistic signals, such as the absence of oscillations (all negative $R^2$ values, denoting poor fit, were set to -1 in the figure). **b**) Estimated parameters (marked as crosses) for each subject, obtained through surrogate optimization, are plotted on the same $\eta$-$\theta$ grid. An orange box highlights the region of the parameter space explored during optimization. This region corresponds to the surrogate optimization interval, within which the parameter sets are considered to best match the subject-specific fMRI data for the 1.5T protocol.

### **Supplementary Text.** Biophysical model

The regional BOLD signals are generated through various transformations in the biophysical model (Sanchez-Rodriguez, Khan, *et al.*, 2024; Sanchez-Rodriguez, Bezgin, *et al.*, 2024). Firstly, the excitatory and inhibitory firing rates in neural mass $k$, denoted as $E_k(t)$ and $I_k(t)$, are derived from the following coupled differential equations:

$$\dot{E}_k = \frac{1}{\tau_E}\left[-E_k + S\left(x_{E,k}\right)\right]$$

$$\dot{I}_k = \frac{1}{\tau_I}\left[-I_k + S\left(x_{I,k}\right)\right]$$

where $x_{E,k} = C_{EE}E_k - C_{IE}I_k + P + \frac{\eta}{N}\sum_{l=1,l\neq k}^{N} \mathrm{C}_{lk}E_l$ and $x_{I,k} = C_{EI}E_k - C_{II}I_k$ are the total input currents received by the excitatory and inhibitory populations, respectively.

The sigmoidal activation functions of the input currents $x_{E,k}$ and $x_{I,k}$ are defined as (Wilson and Cowan, 1972; Daffertshofer and van Wijk, 2011; Gjorgjieva *et al.*, 2016):

$$S_I\left(x_{I,k}\right) = \frac{1}{1 + \exp\left[-a_I\left(x_{I,k} - \theta_I\right)\right]} - \frac{1}{1 + \exp\left[a_I\theta_I\right]}$$

$$S_E\left(x_{E,k}\right) = \frac{1}{1 + \exp\left[-a_E\left(x_{E,k} - \theta_{E,k}\right)\right]} - \frac{1}{1 + \exp\left[a_E\theta_{E,k}\right]}$$

The model's dynamics is modulated by a global scaling factor, ${}^{\eta}/_{N}$ , operating on the cortico-cortical connections, $\mathrm{C}$, where $\eta$ represents coupling strength and $N$ is the total number of modeled brain regions. Additionally, the firing threshold parameters $\theta$ determine the firing properties of the neuronal populations in relation to the local input currents. When $\theta$ is lower than a baseline value, neuronal populations become hyperexcitable, responding to lower levels of input current with an increased firing rate (van Nifterick *et al.*, 2022; Daffertshofer and van Wijk, 2011). Conversely, setting $\theta$ higher than the baseline induces hypoexcitability in neuronal populations. Other parameters are defined in Supplementary Text —Table 1 and are consistent with previous publications (Sanchez-Rodriguez, Khan, *et al.*, 2024; Sanchez-Rodriguez, Bezgin, *et al.*, 2024).

The excitatory/inhibitory (E/I) ratio for the excitatory populations is calculated as the time average of the relative contributions of excitatory and inhibitory synaptic input currents (Soldado-Magraner *et al.*, 2022; Taub *et al.*, 2013; Zhou and Yu, 2018; Xue *et al.*, 2014; Meijer *et al.*, 2015):

$$\langle {}^{E}/_{I} \rangle_k = \left\langle \frac{C_{EE}E_k + \frac{\eta}{N}\sum_{l=1,l\neq k}^{N} \mathrm{C}_{lk}E_l}{C_{IE}I_k} \right\rangle$$

The regional BOLD signal relates to the action potential arriving at the neuronal populations (Logothetis *et al.*, 2001; Sotero and Trujillo-Barreto, 2008; Valdes-Sosa *et al.*, 2009). All quantities are normalized to baseline values:

$$\xi_{E,k} = \frac{s_{E,k}}{{s_{E,k}}^0} \quad \text{(normalized excitatory input)},$$

$$\xi_{I,k} = \frac{s_{I,k}}{{s_{I,k}}^0} \quad \text{(normalized inhibitory input)},$$

where the superscript denotes values at rest.

Changes in glucose consumption ($g_{E,k}$ and $g_{I,k}$) are linked to the excitatory and inhibitory neuronal inputs in region $k$. The glucose variables transform into metabolic rates of oxygen for excitatory ($m_{E,k}$) and inhibitory ($m_{I,k}$) activities, and total oxygen consumption ($m_k$):

$$\dot{g}_{E,k} = z_{E,k}$$

$$\dot{z}_{E,k} = \frac{-2}{\varkappa_E} z_{E,k} - \frac{1}{{\varkappa_E}^2}\left(g_{E,k} - 1\right) + \frac{h_E}{\varkappa_E}\left(\xi_{E,k} - 1\right)$$

$$\dot{g}_{I,k} = z_{I,k}$$

$$\dot{z}_{I,k} = \frac{-2}{\varkappa_I} z_{I,k} - \frac{1}{{\varkappa_I}^2}\left(g_{I,k} - 1\right) + \frac{h_I}{\varkappa_I}\left(\xi_{I,k} - 1\right)$$

$$m_{E,k}(t) = \frac{2 - x(t)}{2 - x_0} g_{E,k}(t)$$

$$m_{I,k}(t) = g_{I,k}(t)$$

$$m_k(t) = \frac{\gamma m_{E,k}(t) + m_{I,k}(t)}{\gamma + 1}$$

$$x(t) = \frac{1}{1 + exp\left[c\left(d - g_{E,k}(t)\right)\right]}$$

Cerebral blood flow ($f_k$) is modeled as follows (Friston *et al.*, 2000), assuming that CBF is coupled to the excitatory activity:

$$\dot{f}_k = y_k$$

$$\dot{y}_k = \frac{-2}{\varkappa_f} y_k - \frac{1}{{\varkappa_f}^2}(f_k - 1) + \mu\left(\xi_{E,k} - 1\right)$$

The outputs of the metabolic and vascular modules are converted to normalized cerebral blood volume ($b_k$) and deoxy-hemoglobin ($q_k$) content through the Balloon model (Buxton *et al.*, 1998):

$$\dot{b}_k = \frac{1}{\varkappa_0}(f_k - f_{out})$$

$$\dot{q}_k = \frac{1}{\varkappa_0}\left(m_k - f_{out}\frac{q_k}{b_k}\right)$$

$$f_{out} = b_k^{\frac{1}{\zeta}}$$

The regional BOLD signal is finally obtained by using the following linear observation equation:

$$BOLD_k(t) = V_0\big(a_1(1 - q_k) - a_2(1 - b_k)\big)$$

where $a_1 = 4.3Y_0E_0 \cdot TE + \varepsilon r_0 E_0 \cdot TE$ and $a_2 = \varepsilon r_0 E_0 \cdot TE + \varepsilon - 1$ are parameters that depend on the experimental conditions (field strength, $TE$) (Obata *et al.*, 2004; Simon and Buxton, 2015; Archila-Meléndez *et al.*, 2020; Deco *et al.*, 2018).

**Supplementary Text —Table 1.** Parameters of the biophysical model

| Parameter | Definition | Value | Ref. |
|---|---|---|---|
| $\begin{bmatrix} E_0 \\ I_0 \\ g_{E0} \\ z_{E0} \\ g_{I0} \\ z_{I0} \\ f_0 \\ y_0 \\ b_0 \\ q_0 \end{bmatrix}$ | Initial conditions | $\begin{bmatrix} 0.075 \\ 0.01 \\ 1 \\ 0 \\ 1 \\ 0 \\ 1 \\ 0 \\ 1 \\ 1 \end{bmatrix}$ | (Sotero and Trujillo-Barreto, 2007, 2008; Sotero *et al.*, 2009; Valdes-Sosa *et al.*, 2009) |
| $\tau_I$ | Time-constant controlling the decay of inhibitory activity after stimulation | 0.02s | (Abeysuriya *et al.*, 2018) |
| $\tau_E$ | Time-constant controlling the decay of excitatory activity after stimulation | 0.01s | (Abeysuriya *et al.*, 2018) |
| $C_{II}$ | Local inhibitory-inhibitory connection strength | 1.2 | (Gjorgjieva *et al.*, 2016; Meijer *et al.*, 2015; Wilson and Cowan, 1972) |
| $C_{EI}$ | Local excitatory-inhibitory connection strength | 6 | (Gjorgjieva *et al.*, 2016; Meijer *et al.*, 2015; Wilson and Cowan, 1972) |
| $C_{EE}$ | Local excitatory-excitatory connection strength | 6.4 | (Gjorgjieva *et al.*, 2016; Meijer *et al.*, 2015; Wilson and Cowan, 1972) |
| $C_{IE}$ | Local inhibitory-excitatory connection strength | 4.8 | (Gjorgjieva *et al.*, 2016; Meijer *et al.*, 2015; Wilson and Cowan, 1972) |
| $P$ | Average constant external input received by the excitatory population | 0.65 | (Gjorgjieva *et al.*, 2016; Meijer *et al.*, 2015; Wilson and Cowan, 1972) |
| $a_I$ | Maximum slope of the inhibitory sigmoidal activation function | 1 | (Abeysuriya *et al.*, 2018) |
| $a_E$ | Maximum slope of the excitatory sigmoidal activation function | 1 | (Abeysuriya *et al.*, 2018) |

| | | | |
|---|---|---|---|
| $\theta_I$ | Position of the inhibitory sigmoidal firing function's threshold for activation | 4 | (Gjorgjieva *et al.*, 2016; Meijer *et al.*, 2015; Wilson and Cowan, 1972) |
| $\theta_E$ | Position of the excitatory sigmoidal firing function's threshold for activation | Variable | (Gjorgjieva *et al.*, 2016; Meijer *et al.*, 2015; Wilson and Cowan, 1972; Daffertshofer and van Wijk, 2011; Abeysuriya *et al.*, 2018) |
| $\eta$ | Global coupling strength scaling the anatomical connectivity matrix $C_{lk}$ | Variable | (Gjorgjieva *et al.*, 2016; Meijer *et al.*, 2015; Wilson and Cowan, 1972; Daffertshofer and van Wijk, 2011; Abeysuriya *et al.*, 2018) |
| $N$ | Number of brain regions of interest | 104 | (Klein and Tourville, 2012; Lin *et al.*, 2022) |
| $h_E$ | Efficacy of glucose consumption response to excitation | 1 | (Sotero and Trujillo-Barreto, 2007, 2008; Sotero *et al.*, 2009; Valdes-Sosa *et al.*, 2009) |
| $h_I$ | Efficacy of glucose consumption response to inhibition | 1 | (Sotero and Trujillo-Barreto, 2007, 2008; Sotero *et al.*, 2009; Valdes-Sosa *et al.*, 2009) |
| $\varkappa_E$ | Time-constant of the excitatory glucose consumption impulse response. | 1s | (Sotero and Trujillo-Barreto, 2007, 2008; Sotero *et al.*, 2009; Valdes-Sosa *et al.*, 2009) |
| $\varkappa_I$ | Time-constant of the inhibitory glucose consumption impulse response. | 1s | (Sotero and Trujillo-Barreto, 2007, 2008; Sotero *et al.*, 2009; Valdes-Sosa *et al.*, 2009) |
| $c$ | Steepness of the sigmoid function $x$ | 2.5 | (Sotero and Trujillo-Barreto, 2007, 2008; Sotero *et al.*, 2009; Valdes-Sosa *et al.*, 2009) |
| $d$ | Position of the threshold of the sigmoid function $x$ | 1.6 | (Sotero and Trujillo-Barreto, 2007, 2008; Sotero *et al.*, 2009; Valdes-Sosa *et al.*, 2009) |
| $\gamma$ | Baseline ratio of excitatory to inhibitory synaptic activity in the voxel | 5 | (Sotero and Trujillo-Barreto, 2007, 2008; Sotero *et al.*, 2009; Valdes-Sosa *et al.*, 2009) |

| | | | |
|---|---|---|---|
| $x_0$ | Fraction of glucose following the glycogenolitic pathway at rest | $\frac{1}{1+exp\left[c\left(d-1(t)\right)\right]}$ | (Sotero and Trujillo-Barreto, 2007, 2008; Sotero *et al.*, 2009; Valdes-Sosa *et al.*, 2009) |
| $\mu$ | Efficacy of blood flow response to excitation | 0.8 | (Sotero and Trujillo-Barreto, 2007, 2008; Sotero *et al.*, 2009; Valdes-Sosa *et al.*, 2009) |
| $\varkappa_f$ | Time constant for CBF response | 1.7 | (Sotero and Trujillo-Barreto, 2007, 2008; Sotero *et al.*, 2009; Valdes-Sosa *et al.*, 2009) |
| $\varkappa_0$ | Transit time through the balloon | 1 | (Sotero and Trujillo-Barreto, 2007, 2008; Sotero *et al.*, 2009; Valdes-Sosa *et al.*, 2009) |
| $\zeta$ | Coefficient of the steady state flow-volume relationship | 0.4 | (Sotero and Trujillo-Barreto, 2007, 2008; Sotero *et al.*, 2009; Valdes-Sosa *et al.*, 2009) |
| $V_0$ | Baseline blood volume | 0.03 | (Sotero and Trujillo-Barreto, 2007, 2008; Sotero *et al.*, 2009; Valdes-Sosa *et al.*, 2009) |
| $E_0$ | Baseline oxygen extraction fraction | 0.4 | (Obata *et al.*, 2004; Simon and Buxton, 2015; Archila-Meléndez *et al.*, 2020) |
| $\Upsilon_0$ | frequency offset of a fully deoxygenated blood vessel | 80.6$s^{-1}$ at 3 T<br>40.3$s^{-1}$ at 1.5 T | (Obata *et al.*, 2004; Simon and Buxton, 2015; Archila-Meléndez *et al.*, 2020) |
| $r_0$ | Slope defining the dependence of the R2* relaxation rate on blood oxygenation | 178$s^{-1}$ at 3 T<br>25$s^{-1}$ at 1.5 T | (Obata *et al.*, 2004; Simon and Buxton, 2015; Archila-Meléndez *et al.*, 2020) |
| $\varepsilon$ | Intrinsic ratio of blood to tissue signals at rest | 0.24 at 3 T<br>1.43 at 1.5 T | (Obata *et al.*, 2004; Simon and Buxton, 2015; Archila-Meléndez *et al.*, 2020) |
| $TE$ | Echo time | 0.030s (3T protocol)<br>0.033s (1.5T protocol) | (Ng *et al.*, 2024) |
| $TR$ | Time to repeat | 3s (3T protocol)<br>2s (1.5T protocol) | (Ng *et al.*, 2024) |
| | Number of volumes acquired | 160 (3T protocol)<br>239 (1.5T protocol) | (Ng *et al.*, 2024) |
